# 24-Nor-Ursodeoxycholic acid reshapes immunometabolism in CD8^+^ T cells and alleviates hepatic inflammation

**DOI:** 10.1101/2021.01.09.426037

**Authors:** Ci Zhu, Nicole Boucheron, André C. Müller, Peter Májek, Thierry Claudel, Emina Halilbasic, Hatoon Baazim, Alexander Lercher, Csilla Viczenczova, Daniela Hainberger, Teresa Preglej, Lisa Sandner, Marlis Alteneder, Alexandra F. Gülich, Matarr Khan, Patricia Hamminger, Jelena Remetic, Anna Ohradanova-Repic, Philipp Schatzlmaier, Clemens Donner, Claudia D. Fuchs, Tatjana Stojakovic, Hubert Scharnagl, Shinya Sakaguchi, Thomas Weichhart, Andreas Bergthaler, Hannes Stockinger, Wilfried Ellmeier, Michael Trauner

## Abstract

**Background & Aims:** 24-NorUrsodeoxycholic acid (NorUDCA) is novel therapy for immune-mediated liver diseases such as primary sclerosing cholangitis (PSC) where dysregulated T cells including CD8^+^ T cells cause liver immunopathology. We hypothesized that NorUDCA may directly modulate CD8^+^ T cell effector function thus contributing to its therapeutic efficacy independent of anti-cholestatic effects.

**Methods:** NorUDCA effects on CD8^+^ T cell function *in vivo* were investigated in a hepatic injury model system induced by excessive CD8^+^ T cell immune response upon non-cytolytic lymphocytic choriomeningitis virus (LCMV) infection. Mechanistic studies included molecular and biochemical approaches, flow cytometry and metabolic assays in mouse CD8^+^ T cells *in vitro*. Mass spectrometry (MS) was used to identify potential targets modulated by NorUDCA in CD8^+^ T cells. NorUDCA signaling effects observed in murine systems were validated in peripheral T cells from healthy volunteers and PSC patients.

**Results:** *In vivo* NorUDCA ameliorated hepatic injury and systemic inflammation upon LCMV infection. Mechanistically, NorUDCA demonstrated a strong immunomodulatory efficacy in CD8^+^ T cells affecting lymphoblastogenesis, mTORC1 signaling and glycolysis of CD8^+^ T cells. With MS, we identified that NorUDCA regulates CD8^+^ T cells via targeting mTORC1. NorUDCA’s impact on mTORC1 signaling was further confirmed in circulating human CD8^+^ T cells.

**Conclusions:** NorUDCA possesses a yet-unrecognized direct modulatory potency on CD8^+^ T cells and attenuates excessive CD8^+^ T cell hepatic immunopathology. These findings may be relevant for treatment of immune-mediated liver diseases such as PSC and beyond.

## Introduction

CD8^+^ T cells are critical players of cell-mediated adaptive immunity and provide key defense mechanisms against microbial infections and cancer^1^. However, when CD8^+^ T cell immune responses and tissue infiltration become dysregulated and excessive, CD8^+^ T cells can turn into immunopathological factors driving hepatic inflammation and damage^2^. As such, CD8 T cell-mediated cytotoxicity and tissue damage play important role in viral, autoimmune or immune-mediated liver diseases^3, 4^ Therefore identifying novel therapies to control detrimental CD8^+^ T cell inflammatory reactions is pivotal to counteract the increasing burden of autoimmune liver diseases and the corresponding need for liver transplantation, reflecting the limited effectiveness of current therapeutic options^5^

24-NorUrsodeoxycholic acid (NorUDCA) is a novel therapeutic bile acid (BA) whose pharmacological mechanisms profoundly differ from its biochemical parent compound ursodeoxycholic acid (UDCA) which has been used with variable success in a wide range of cholestatic and metabolic liver diseases^6^. In a recent phase II study, NorUDCA has demonstrated promising results in treating primary sclerosing cholangitis (PSC)^7^, an immune-mediated cholestatic liver disease so far lacking effective medical therapy. Dysregulated T cells including CD8^+^ T cells play a critical role in mediating immunopathogenesis of PSC^4^. Therefore we hypothesized that NorUDCA’s therapeutic mechanism of action may be attributed to potential immunomodulatory potency on CD8^+^ T cells besides its well-established anti-cholestatic efficacy^8^.

We first explored NorUDCA’s impact on CD8^+^ T cell effector function *in vivo* in a non-cholestatic murine model based on the non-cytolytic lymphocytic choriomeningitis virus (LCMV) infection, in which hepatic injury and systematic inflammation are predominantly mediated by excessive cytotoxicity of effector CD8 T cells at early phase of infection^9-11^. Further mechanistic studies were performed using molecular and biochemical approaches, flow cytometry and metabolic assays in primary murine CD8^+^ T cells *in vitro*. Mass spectrometry (MS) was used to identify potential targets modulated by NorUDCA in CD8^+^ T cells. Human peripheral T cells from healthy volunteers and PSC patients were used to validate NorUDCA signaling effects observed in murine systems.

## Results

### NorUDCA reduces frequency and cell size of intrahepatic effector CD8^+^ T cells with alleviated hepatic injury *in vivo*

First we explored the effect of NorUDCA on effector CD8^+^ T cell immune response in a murine model induced by LCMV Clone 13 infection specifically focusing on the acute phase of the infection on day 10 when escalation of hepatic injury had been widely reported^9-11^, a disease phenotype being absent in LCMV Armstrong strain^10^. This allows us to assess if NorUDCA impacts on effector CD8^+^ T cell response causing hepatic injury in absence of cholestasis. NorUDCA’s parent compound UDCA was used in this model for comparison.

In line with previous reports^9, 11-13^, we observed a significant loss of body weight and elevated level of hepatic injury parameters such as ALT, AST and AP in mice following LCMV infection, which were significantly attenuated by NorUDCA, while UDCA showed only minimal effects (Fig. 1A, 1B).

We observed that the frequency of intrahepatic CD8^+^ T cells positive for the major LCMV epitope glycoprotein-33 (GP33) in NorUDCA-treated infected mice was lower compared with untreated UDCA-treated infected mice (Fig.1C, gating strategy see Supplementary Fig. 1). Despite the change of frequency of intrahepatic virus-specific CD8^+^ T cells in NorUDCA-treated infected mice, the effector function of virus-specific CD8^+^ T cells remained unchanged, as evidenced by the comparable expression of cytolytic molecules of Granzyme B and Perforin in intrahepatic virus-specific effector CD8^+^ T cells in infected mice treated with and without NorUDCA (Fig. 1D). This might explain why the hepatic viral loads were not affected by NorUDCA treatment in infected mice even though less intrahepatic CD8^+^ T cells were present (Fig. 1E).

**Fig. 1.**
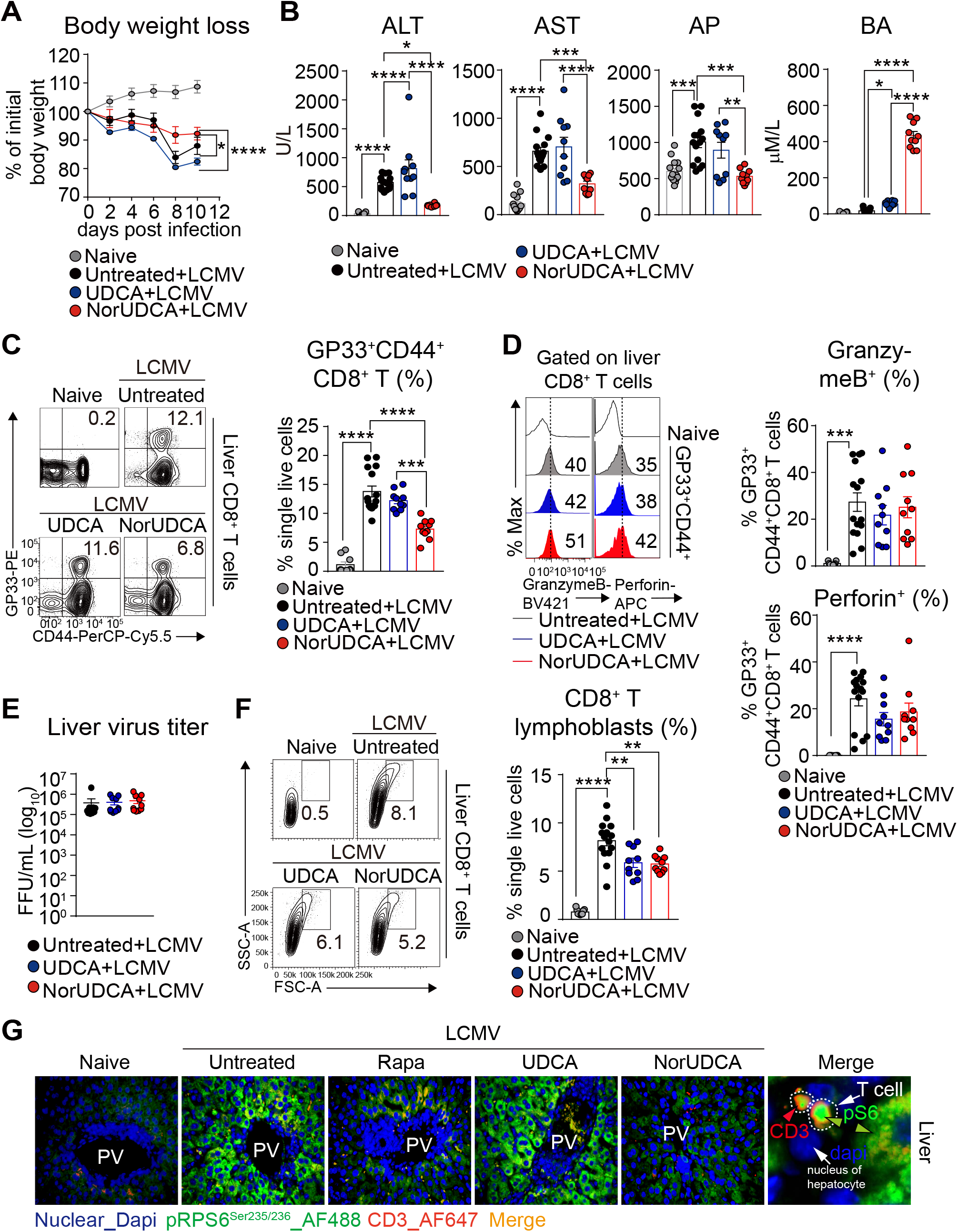
NorUDCA suppresses cell size and frequency of intrahepatic effector CD8^+^ T cells in LCMV model. (A) Changes in body weight over time. (B) Serum liver enzymes ALT, AST, AP, BA. (C) Representative plots and quantitative analysis of intrahepatic virus-specific CD8^+^ T cells. (D) Representative plots of intrahepatic virus-specific CD8^+^ expressing cytolytic molecules. Quantitative analysis is shown alongside. (E) Liver LCMV virus titer. (F) Representative plots and quantitative analysis of intrahepatic CD8^+^ T lymphoblasts. (G) Representative high power field multicolor-immunofluorescence staining of CD3 and pRPS6^Ser235/236^ of liver slides of indicated groups. Data are summary of 3 independent experiments. At least 3 biologically independent animals were used per group during experiments. Quantitative data are presented as mean±SE. *P* values were calculated by one-way ANOVA corrected with Tukey post-hoc test and in A were calculated by two way ANOVA. *=*P*<0.05, **=*P*<0.01, ***=*P*<0.001, ***=*P*<0.001, ****=*P*<0.0001. ALT, alanine aminotransferase; AST, aspartate transaminase; AP, alkaline phosphatase; BA, bile acid; PV, portal vein; Rapa, Rapamycin; Dapi, 4’, 6-diamidino-2-phenylindole; AF, Alexa Fluor.

Of note, we observed that the cell size of intrahepatic CD8^+^ T cells isolated from LCMV mice treated with NorUDCA was much smaller compared to that of untreated LCMV mice (Fig. 1F). Since a large number of studies indicate that cell size of CD8^+^ T cells are critically controlled by mTORC1 signaling^14, 15^, we next investigated if mTORC1 signaling in intrahepatic CD8^+^ T cells is affected in NorUDCA-treated infected mice. Therefore, for the subsequent analysis of mTORC1 signaling, we included Rapamycin as mTORC1 inhibitor control (at a concentration used in a previous LCMV infection study^16^). We performed multicolor immunofluorescence microscopy of the liver sections to assess the phosphorylation of Ribosomal protein S6 (RPS6) RPS6^Ser235/236^ (a direct mTORC1 downstream target) as read out for mTORC1 activity. We detected an increase of hepatic infiltrating pRPS6^Ser235/236^-positive CD3^+^ T cells in mice upon LCMV infection, which were decreased when treated with Rapamycin or NorUDCA (Fig. 1G). This suggests that mTORC1 is repressed by Rapamycin and NorUDCA. In addition we also detected a downregulation of pRPS6^Ser235/236^ in hepatocytes from Rapamycin-or NorUDCA-treated infected mice (Fig. 1G), suggesting that the inhibitory efficacy of Rapamycin and NorUDCA is not limited to CD8^+^ T cells but also expands to other cell types. In contrast, pRPS6^Ser235/236^ levels were not altered in UDCA-treated LCMV mice, indicating that UDCA does not impact mTORC1 signaling in LCMV infected mice (Fig. 1G). In summary, our results indicate that NorUDCA treatment during acute phase of LCMV infection led to a reduction in effector CD8^+^ T cell-driven inflammation and injury.

### NorUDCA modulates effector CD8^+^ T cells systemically *in vivo*

To determine whether the observed immunomodulatory activity of NorUDCA observed in LCMV model is restricted to liver or can be extended to extrahepatic tissues, we analyzed NorUDCA’s impacts on the CD8^+^ T cells from blood and spleen. Similarly to our observation made in liver, NorUDCA reduced frequency of virus-specific effector CD8^+^ T cells in both blood and spleen in infected mice, while UDCA had no impact (Fig. 2A). Further we found that, like Rapamycin, NorUDCA decreased the phosphorylation (pSer235/236) of mTORC1 downstream target RPS6 in circulating effector CD8^+^ T cells of infected mice (Fig. 2B).

**Fig. 2.**
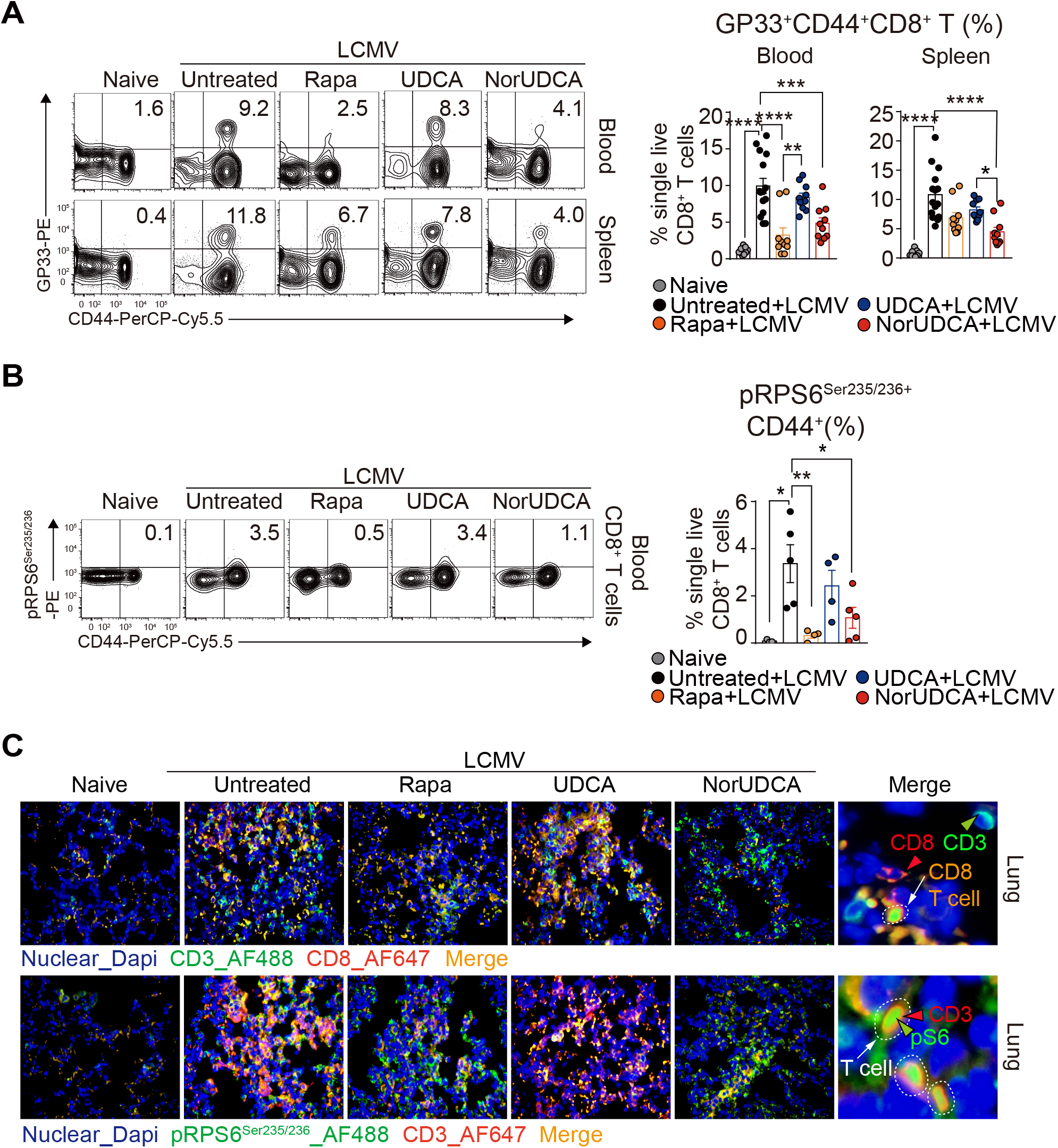
NorUDCA exerts strong systemic immunometabolic modulatory properties *in vivo*. (A) Representative plots of virus-specific CD8^+^ T cells from peripheral blood and spleen. Quantitative analysis is shown alongside. (B) pRPS6^Ser235/236^ expression on blood CD8^+^ T cells. (d) Quantitative analysis of is shown alongside. (C) Representative high power field multicolor-immunofluorescence staining of CD3 and pRPS6^Ser235/236^ (upper) and CD3 and CD8 of lung slides of indicated groups. Data are pooled from 3 independent experiments. At least 3 biologically independent animals were used per group during experiments. Quantitative data are presented as mean±SE. *P* values were calculated by one-way ANOVA corrected with Tukey post-hoc test. *=*P*<0.05, **=*P*<0.01, ***=*P*<0.001, ***=*P*<0.001, ****=*P*<0.0001.

Moreover, we also investigated if NorUDCA impacts on mTORC1 signaling in splenic effector CD8^+^ T cells *ex vivo*. To mimic the *in vivo* activation, we isolated splenocytes from infected mice treated with or without NorUDCA, UDCA or Rapamycin and re-stimulated the cells with LCMV-GP33 peptides. pRPS6^Ser235/236^ was assessed in effector CD8^+^ T cells (Supplementary Fig. 2A). pRPS6^Ser235/236^ levels were lower in the splenic effector CD8^+^ T cells displaying surface expression of CD107a/b (a degranulation marker indicative of effector CD8^+^ T cell function) in NorUDCA-treated infected mice upon re-stimulation showing a reduced mTORC1 activity (Supplementary Fig. 2B).

Of note, upon *ex vivo* re-stimulation, splenic effector CD8^+^ T cells from NorUDCA-treated infected mice expressed reduced levels of TNFα. However, IFNγ expression, which is essential for maintaining efficient protective immunity against LCMV^17^, was comparable (Supplementary Fig. 2C) indicating a potential intact anti-viral immunity. This was further supported by the unchanged splenic virus titer in NorUDCA treated-infected mice (Supplementary Fig. 2D).

Additionally, we analyzed CD8^+^ T cell infiltration in the lung as another extrahepatic organ. Multicolor immunofluorescence microscopy revealed that NorUDCA not only reduced infiltrating CD8^+^ T cells but also T cells displaying pRPS6^Ser235/236^ in the lung parenchyma of infected mice, further suggesting a systemic modulatory effect of NorUDCA on mTORC1 during CD8^+^ T cell differentiation *in vivo* (Fig. 2C).

### NorUDCA reduces CD8^+^ T cell blasting and expansion associated with attenuated mTORC1 signaling *in vitro*

To gain further mechanistic insights about NorUDCA’s direct impact on CD8^+^ T cell signaling, function and metabolism (see Supplementary Fig. 3 for gating strategy) we activated primary murine CD8^+^ T cells from spleen and lymph nodes *in vitro* in the presence of NorUDCA or UDCA with the concentrations precisely matching the *in vivo* concentrations in our LCMV model system (Fig. 1B).

Upon antigen recognition, naïve CD8^+^ T cells undergo a complex activation, blasting (increasing cell size), proliferation and differentiation process leading to establishment of a large pool of various effector CD8^+^ T cells required to mount effective immunity^1^. Both NorUDCA and UDCA showed no effects on T cell receptor-proximal CD8^+^ T cell activation as reflected by unchanged 9. up-regulation of T cell activation markers CD25 and CD69 and normal Ca^2+^ influx (Fig. 3A, 3B). However, CD8^+^ T cell blasting (Fig. 3C) and clonal expansion (Fig. 3D) were significantly reduced by NorUDCA without affecting cell viability (Fig. 3C) compared to cells without treatment or treated with UDCA. In accordance to our *in vivo* mTORC1 signaling finding, NorUDCA decreased the frequency of CD8^+^ T cells displaying high level of pRPS6^S235/236^ (Fig. 3E), indicating that NorUDCA impairs mTORC1 activity by acting directly on CD8^+^ T cells during their differentiation to effector cells.

**Fig. 3.**
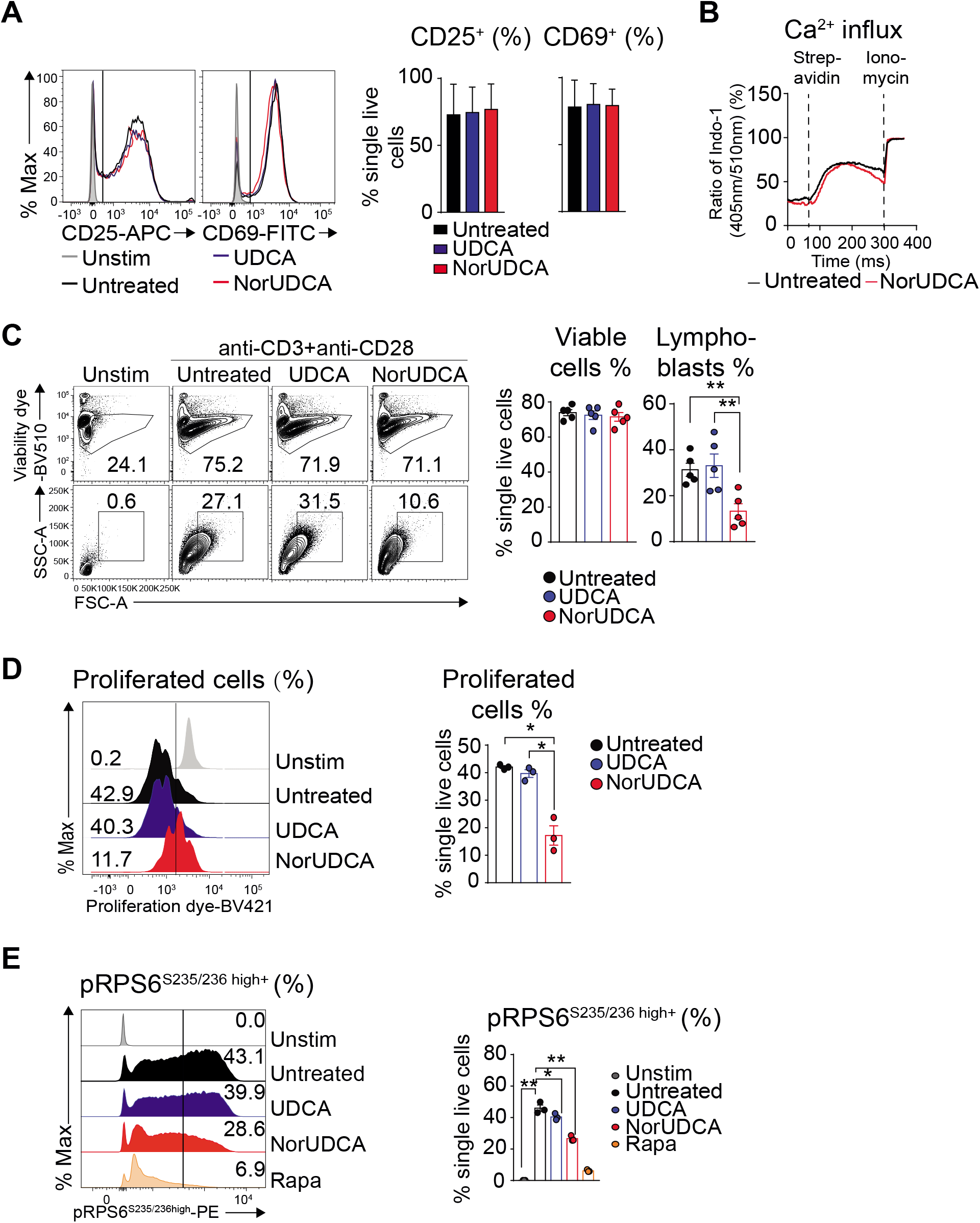
NorUDCA attenuates murine CD8^+^ T cells blasting and expansion with blunted mTORC1 signaling. (A) Primary murine CD8^+^ T cells were stimulated for 24h using anti-CD3 and anti-CD28 mAbs ± compound treatment as indicated. Histograms show CD25 and CD69 expression. Quantification is shown alongside. (B) Graph represents calcium influx in activated CD8^+^ T cells ± NorUDCA. (C) Primary murine CD8^+^ T cells were stimulated for 36h under indicated condition. Numbers indicate frequencies of viable cells or lymphoblasts. Quantitative analysis is shown alongside. (D) Proliferation of primary murine CD8^+^ T cells treated as in (C) by dilution of intracellular proliferation dye. Numbers show frequency of proliferated cells. Quantitative analysis is shown alongside. (E) Histograms depict pRPS6^S235/236^ in CD8^+^ T cells treated as in (C). Numbers in the histograms indicate frequencies of cells with high pRPS6^S235/236^ expression. Quantitative analysis is shown alongside. Data are representative of 3 independent experiments. Quantitative data are presented as mean±SE. *P* values were calculated by one-way ANOVA corrected with Tukey post-hoc test. *=*P*<0.05, **=*P*<0.01. Unstim, unstimulated, Stim, stimulated, Rapa, Rapamycin.

### NorUDCA redirects activation-induced metabolic reprograming in CD8^+^ T cells

Since mTORC1 is required for the expression of glucose transporters and enzymes that control glycolysis in CD8 T cells, we therefore explored whether NorUDCA alters the metabolic profile of CD8^+^ T cells. *In vitro* we found that NorUDCA repressed the expression of genes in activated CD8^+^ T cells that promote glycolysis. Surface GLUT1 expression on CD8^+^ T cells was also reduced by NorUDCA (Fig. 4A-C), which is in line with our *in vivo* observation. Furthermore NorUDCA reduced intracellular synthesis and output of lactate in CD8^+^ T cells, an end product of glycolytic pathway (Fig. 4D). To explore whether inhibition of glycolysis enforces fatty acid β-oxidation (FAO) in CD8^+^ T cells ^19^, we tested the expression of genes regulating FAO in CD8^+^ T cells. NorUDCA significantly upregulated carnitine palmitoyltransferase 1a d (Cpt1a) expression (Fig. 4A), a rate-limiting enzyme in FAO

**Fig. 4.**
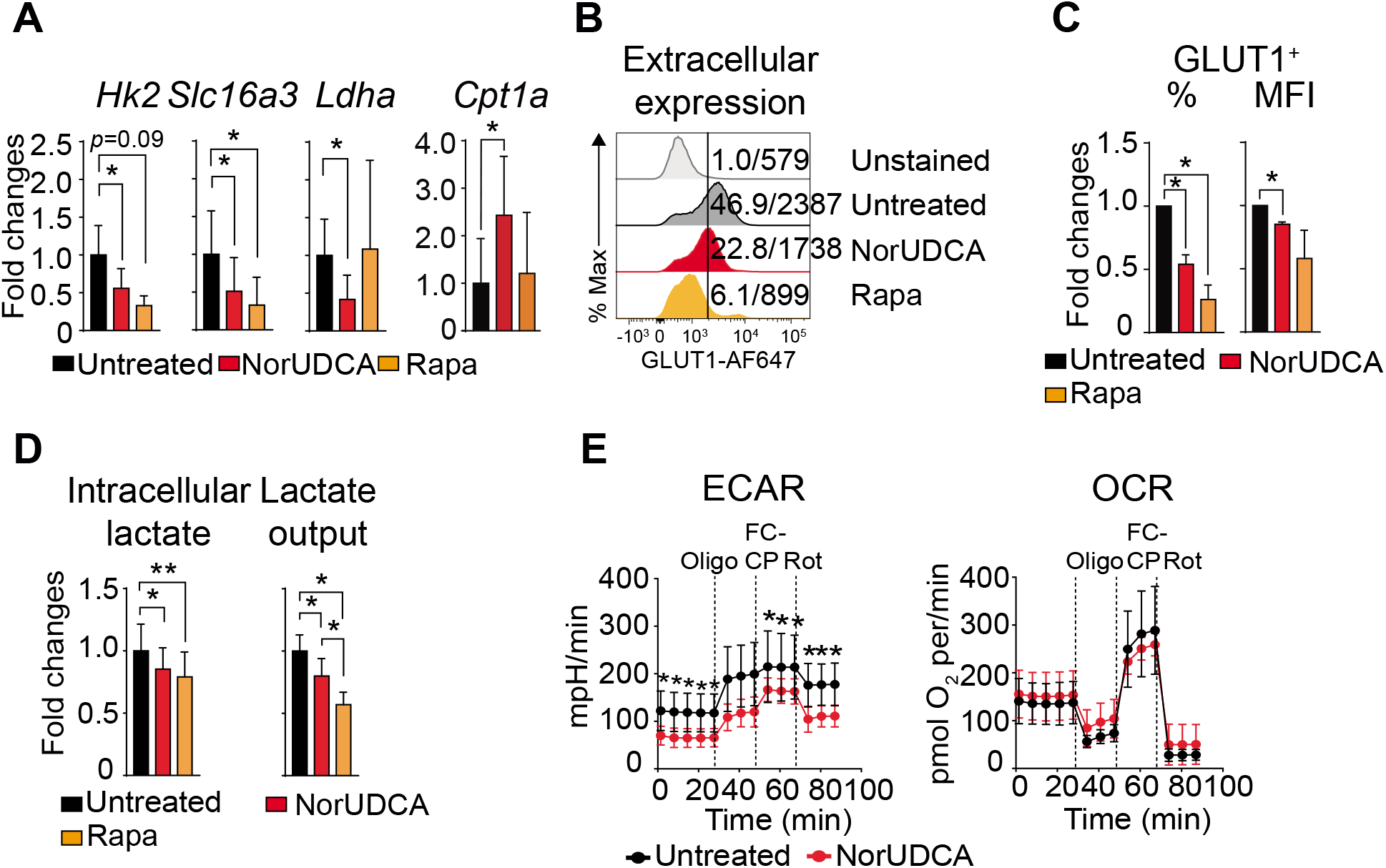
NorUDCA demonstrates immunometabolic actions in modulating murine CD8^+^ T cell *in vitro*. (A) Quantitative RT-qPCR analysis of expression of indicated genes (normalized to *Hprt* and untreated groups) in primary murine CD8^+^ T cells stimulated for 24h ± compound treatments as indicated. (B) Representative histograms depicting GLUT1 extracellular expression of cells cultured as in (A) and treated as indicated. Quantitative analysis is shown alongside. (C) Quantification of intra- and extra-cellular output of lactate from cells cultured as in (A). (D) Real time changes in the extracellular acidification rate (ECAR; top) and oxygen consumption rate (OCR; bottom) of activated CD8^+^ T cells after treatment with oligomycin (Oligo), Carbonyl cyanide-4-(trifluoromethoxy)phenylhydrazone (FCCP), and rotenone (Rot) ± NorUDCA. Data are either representative or show the summary of 3 independent experiments. Summary data are presented as mean±SE. *P* values were calculated by one-way ANOVA corrected with Tukey post-hoc test. *= *P*<0.05, **=*P*<0.01. Rapa, Rapamycin.

To further assess the functional impact of NorUDCA on glycolysis and mitochondria respiratory activity, we performed extracellular metabolic flux measurements to determine real-time changes in extracellular acidification rate (ECAR) and oxygen consumption rate (OCR). NorUDCA decreased the glycolytic activity in CD8^+^ T cells, indicated by a lower level of ECAR, while OCR (indicator for mitochondria oxidative phosphorylation (OXPHOS)) was not affected in this setting (Fig. 4E)

### NorUDCA reshapes mTORC1 associated (phospho-)proteome and metabolic landscape in CD8^+^ T cells upon activation

To mechanistically understand how NorUDCA impact on mTORC1, we performed quantitative high-resolution mass spectrometry (MS) using murine CD8^+^ T cells treated with NorUDCA or Rapamycin. We first focused on comparing NorUDCA with Rapamycin regarding their modulation of mTORC1-associated (phospho-)proteome in CD8^+^ T cells. In total 7,492 proteins and 15,538 phosphorylated sites per biological replicate were identified (Fig. 5A). Cluster analysis of differentially expressed proteins and phosphopeptides demonstrated that NorUDCA had similarities with Rapamycin in reshaping the overall (phospho-)proteome of CD8^+^ T cells (Fig. 5B). Analysis revealed that 80.1% (1,543 of 1,926 proteins) of the proteins significantly modulated by NorUDCA were also modulated by Rapamycin in the same direction, further revealing mTORC1 as a main pathway targeted by NorUDCA. Interestingly we noticed that still 19.1% (383 of 1,926 proteins) were uniquely modulated by NorUDCA (Fig. 5C). Further functional annotations by Hallmark, GO (gene ontology) and KEGG databases identified metabolic pathways shared by NorUDCA and Rapamycin, as well as pathways uniquely affected by NorUDCA (Supplementary Fig. 4, Supplementary Table 1).

**Fig. 5.**
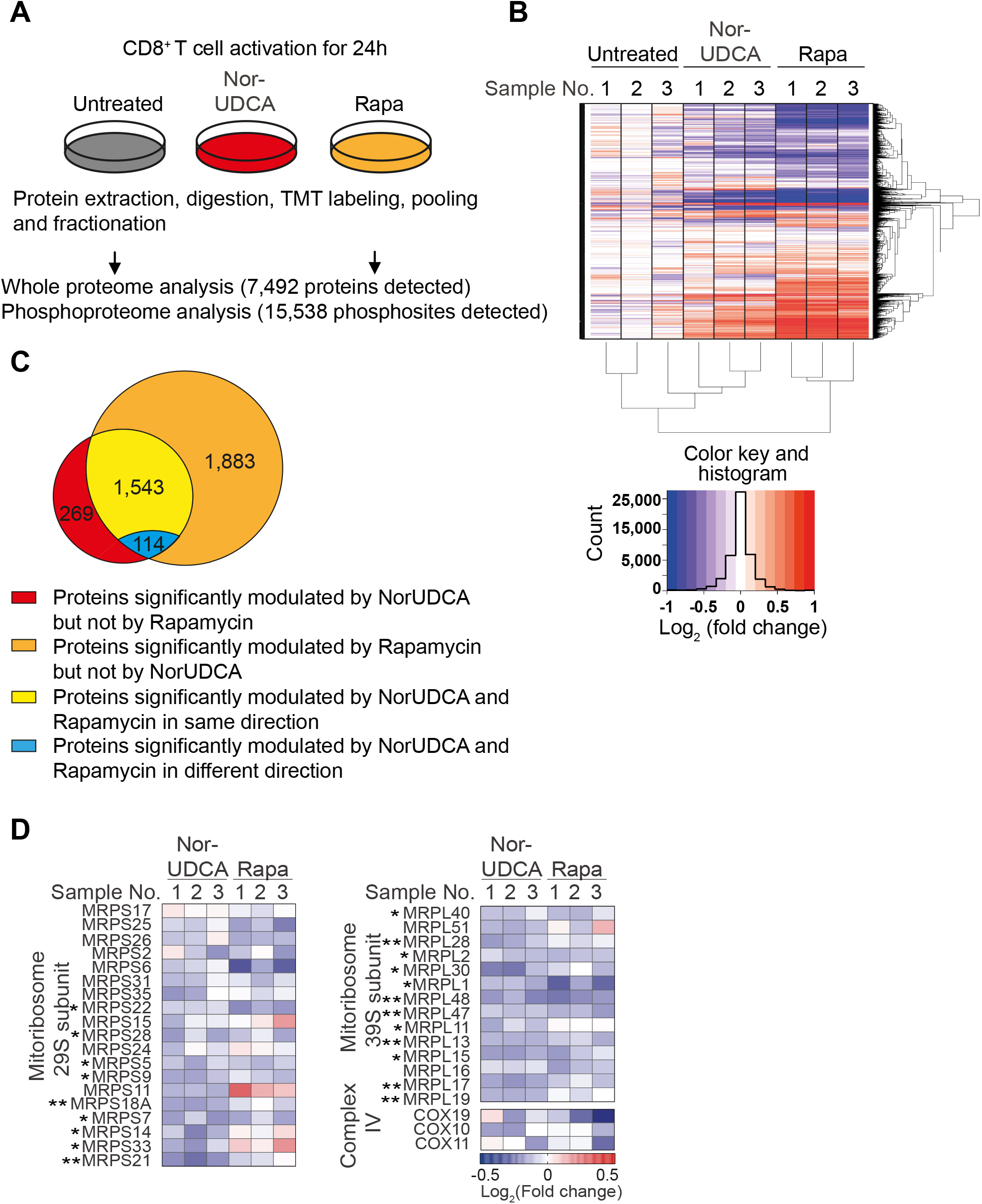
NorUDCA shows similar impacts on CD8^+^ T cell proteomic profile and reduces mitoribosome protein abundance in CD8^+^ T cells like Rapamycin. (A) Scheme depicting the experimental approach. (B) Hierarchical cluster analysis of differentially regulated proteins under indicated conditions are shown in the heat map represented color-coded expression levels following Log_2_ value of ratio between protein content of NorUDCA-or Rapamycin-treated CD8^+^ T cells and that of untreated cells (n=3). (C) Venn diagrams display the overlap of identified proteins between NorUDCA and Rapamycin treated groups. (D) Heat map of mitoribosome protein abundance changes upon NorUDCA or Rapamycin treatment (both normalized to untreated group) of CD8^+^ T cells cultured as in (A). Data were obtained from proteomics analysis using biological replicates from 3 independent experiments. Asterisks next to protein labeling represent significance of comparison between NorUDCA and untreated groups. Data show the summary of 3 independent experiments. *P* values in were calculated by one-way ANOVA corrected with Tukey post-hoc test. *P* values in (D) were calculated among the comparison between untreated and NorUDCA by two-way Anova. *= *P*<0.05, **=*P*<0.01. Rapa, Rapamycin.

We next studied mTORC1-associated metabolic proteomes modulated by NorUDCA in CD8^+^ T cells. In agreement with our findings described above, we detected a reduction of Hexokinase 2 and additional proteins crucial for glycolysis in MS data set of NorUDCA-treated CD8^+^ T cells (Supplementary Fig.5). Moreover, NorUDCA enhanced protein abundance of FA transporters and enzymes, such as, Long-chain FA transport protein 4 (SLC27A4), Fatty-acyl-CoA synthase (FACS), CPT1α and CPT2 (Supplementary Fig. 5). In contrast, FA synthesis was reduced as revealed by decreased abundance of FA synthase and increased phosphorylation of acetyl-CoA carboxylase (ACACA) at S117 residue whose phosphorylation inhibits enzyme activity (Supplementary Fig. 5). Taken together, these data indicate that NorUDCA redirects CD8^+^ T cell nutrient sensing program from glycolysis toward fatty acid utilization upon activation as consequence of mTORC1 suppression.

Interestingly, MS analysis revealed that the abundance of several mTORC1-dependent mitoribosomes was reduced in NorUDCA-treated CD8^+^ T cells (Fig. 5D). Additionally, mTORC1-regulated COX10 ^20^, which is important for expression and assembly of electron transport components (ETC) complex IV, also showed a tendency for reduction by NorUDCA (Fig. 5D). Overall, our MS analysis revealed a modulation of mTORC1-associated metabolism in CD8^+^ T cells by NorUDCA.

The activity of mTORC1 is regulated by a complex regulatory network. When we tested potential candidate upstream signaling molecules regulating mTORC1, we found that Ras-Erk-P90 RSK signaling was suppressed by NorUDCA, while classic mTORC1 upstream axes involving phosphatidylinositol-3 kinase (PI3K) and AKT^T308^ were not reduced (Supplementary Fig. 6A-C). Additionally intracellular phosphatidic acid (PA) that can induce direct mTORC1 activation by competing with FK506-binding protein 38 for binding with FK506 protein, a well-known endogenous inhibitor of mTORC1 ^21^, was reduced by NorUDCA in activated CD8^+^ T cells (Supplementary Fig. 6D). Collectively, our findings demonstrate that mTORC1 is a central hub for NorUDCA’s signaling action and that NorUDCA affects mTORC1 directly and its upstream activation pathways.

### NorUDCA suppresses human CD8^+^ T cell blasting and expansion with blunted mTORC1 kinase activity

Finally we investigated whether some of our key findings obtained with murine cells apply to human peripheral CD8^+^ T cells, especially peripheral CD8^+^ T cells from patients of PSC, where dysregulated T cells are associated with disease progression^4^. We examined the impact of NorUDCA on CD8^+^ T cell activation, blasting, clonal expansion and mTORC1 *in vitro* by activating *ex vivo* isolated peripheral T cells from healthy volunteers and PSC patients in the absence or presence of NorUDCA (Fig. 6, gating strategy see Supplementary Fig. 7). UDCA was used for comparison. Rapamycin was used as control for mTORC1 inhibition.

**Fig. 6.**
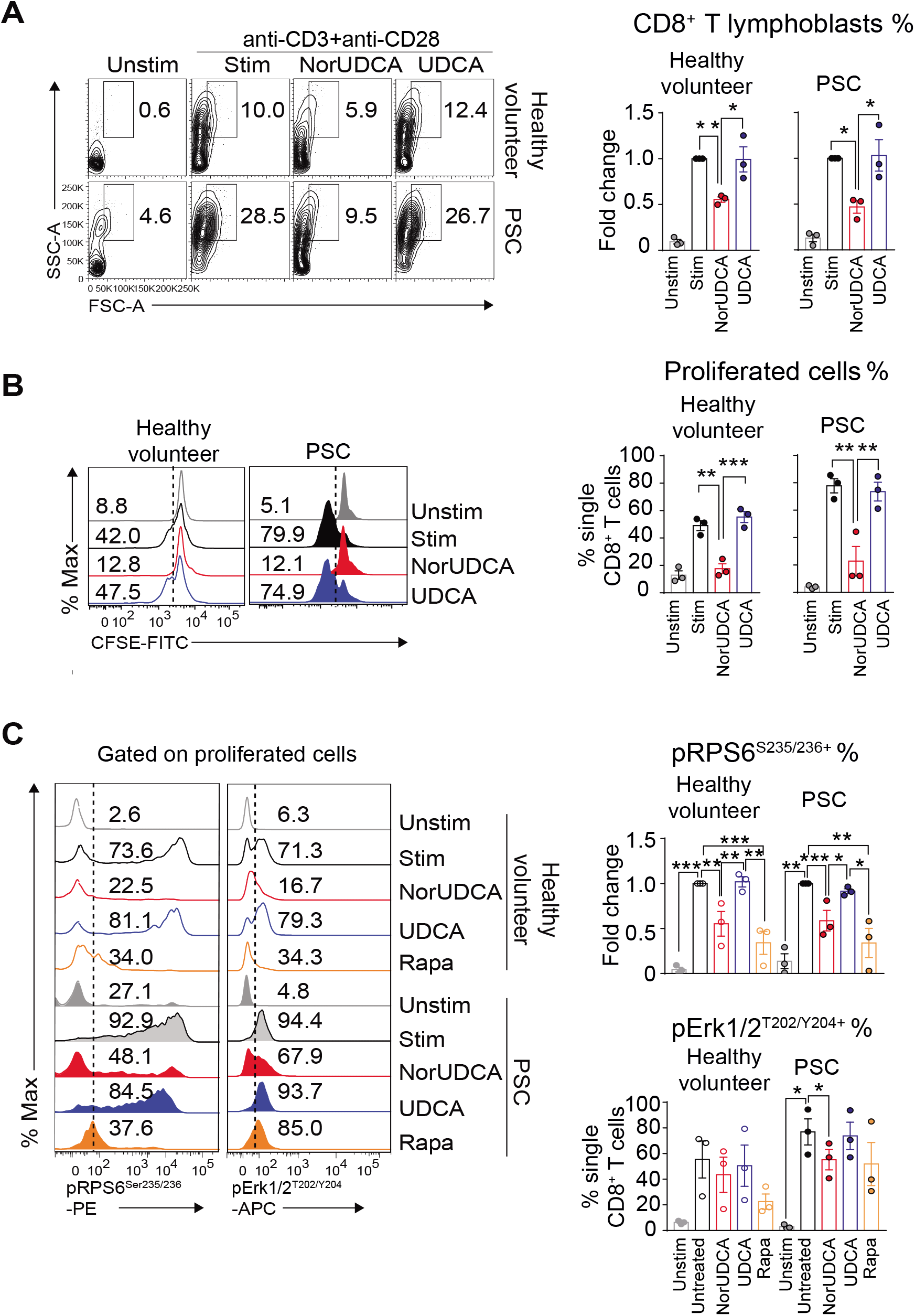
NorUDCA reduces human CD8^+^ T cells lymphoblastogenesis and proliferation with suppressed mTORC1 kinase activities. (A) Peripheral human T cells from age and gender matched healthy volunteers and primary sclerosing cholangitis (PSC) patients were activated under indicated conditions. Representative plots showing lymphoblastogenesis on gated CD8^+^ T cells under indicated conditions. Numbers indicate frequencies of lymphoblasts. Quantitative analysis is shown alongside. (B) Plots display proliferation. Quantitative analysis is shown alongside. (C) pErk1/2^T202/Y204^ and pRPS6^Ser235/236^ on proliferated human CD8^+^ T cells as indicated. Numbers show frequency of positive cells. Quantitative analysis is shown alongside. Data are summary of 3 independent experiments. Quantitative data are presented as mean±SE. *P* values were calculated by one-way ANOVA corrected with Tukey post-hoc test. *=*P*<0.05, **=*P*<0.01, ***=*P*<0.001, ***=*P*<0.001. Unstim, unstimulated, Stim, stimulated, PSC, primary sclerosing cholangitis, Rapa, Rapamycin.

In line with our observations gained from the murine experiments, NorUDCA strongly decreased cell size and clonal expansion in CD8^+^ T cells from both PSC patients and healthy volunteers, while not affecting cell viability and activation. We next explored if NorUDCA affects Erk-mTORC1 signaling in human CD8^+^ T cells by assessing expression of pErk1/2^T202/Y204^ and pRPS6^Ser235/236^. After NorUDCA treatment, we detected a reduced Erk-mTORC1 signaling in proliferating CD8^+^ T cells of healthy volunteers and PSC patients as reflected by reduced frequency of pErk1/2^T202/Y204+ and^ pRPS6^Ser235/236+^ cells, while UDCA showed no effect (Fig. 6C).

## Discussion

The therapeutic mechanisms of NorUDCA as novel therapeutic approach to immune-mediated liver diseases such as PSC^7^ are currently of great interest to the scientific community. Previously we have reported that NorUDCA strongly reversed cholestasis and biliary fibrosis in Mdr2 KO model of sclerosing cholangiopathy with pronounced repression of hepatic inflammation^8^. Since Mdr2 KO mice suffer from defective bile formation (i.e., the lack of biliary phospholipid secretion with consecutive toxic bile formation) and NorUDCA also has substantial effects on bile formation and composition by inducing cholehepatic shunting and subsequently a bicarbonate rich choleresis^22^, it remains unclear whether this promising compound has direct anti-inflammatory effect or the improved hepatic inflammation was indirectly due to NorUDCA’s well-established anti-cholestatic efficacy.

Here we report that NorUDCA is able to exert direct immunomodulatory and anti-inflammatory efficacy in a non-cholestatic immune-mediated model of hepatic and systematic immunopathology upon acute LCMV infection. Given the complexity of how the immune network functions upon infection, we certainly cannot rule out other possible beneficial mechanisms induced by NorUDCA such as modulation of innate and humoral immunity, especially as mTORC1 inhibition was also observed in other cell types than CD8^+^ T cells, namely hepatocytes, we can assume that also other immune cells like antigen presenting cells might be regulated by NorUDCA. However, in this study we uncovered one potential key mechanism by demonstrating that NorUDCA modulates mTORC1 signaling in effector CD8^+^ T cells (a cell type predominantly accounting for the disease phenotype in this model^9-11^) which subsequently influenced on cell frequency, metabolism and function. Whether NorUDCA impacts on antigen presenting cells which mounts subsequent CD8^+^ T cell immune response requires future full-scale study. However, following *in vitro* study and MS analysis, we confirmed NorUDCA has a direct impact on CD8^+^ T cell blasting, proliferation and metabolism and identified mTORC1 as one of the potential targets for its mode of actions, which we believe should advance our mechanistic understandings about this novel compound and extension of future clinical applications.

mTORC1 inhibition provides a critical immunometabolic link controlling CD8^+^ T cell immune response ^18, 23^, regulating glycolysis and FAO ^24^ NorUDCA inhibited glycolysis and enhanced the FAO machinery in CD8^+^ T cells, as reflected by an increased expression of key regulators of FAO. In addition, we observed that OCR of NorUDCA-treated CD8^+^ T cells was unchanged. Mitoribosome plays a critical role in translation of mitochondria-encoded electron transport components to facilitate OXPHOS during T cell quiescence exit ^20^. Our MS data showed that mitoribosome biogenesis and complex IV were disrupted by NorUDCA, which might counteract the up-regulated FAO machinery, and might explain why OCR remained unchanged by NorUDCA.

Although NorUDCA effects share many similarities with classic mTORC1 inhibitor Rapamycin in modulating mTORC1 associated (phospho-)proteome, we detected important differences between the activities of NorUDCA and Rapamycin in modulating CD8^+^ T cell proteomes. Firstly, NorUDCA affects less proteins than Rapamycin. Secondly, an array of metabolic pathways is uniquely altered by NorUDCA, such as metabolism of organic acids, organonitrogen compounds and cellular catabolic processes. Therefore, although the mTORC1 pathway is a potential target of NorUDCA, NorUDCA shows immunometabolic properties that are not exclusively the consequence of a direct mTORC1 inhibition. This indicates exciting entry points for future relevant full-scale metabolic studies.

Notably, UDCA, the structurally closest related clinically used bile acid^6^ and parent compound of NorUDCA, entirely lacked the signaling effects and immunometabolic modulatory functions throughout our comprehensive study comparing NorUDCA and UDCA *in vivo* and *in vitro*. When we tested if our signaling finding could be translated into circulating CD8^+^ T cells of PSC patients, we found that NorUDCA, but not UDCA, suppressed mTORC1 signaling following the concentrations achieved *in vivo* in murine model system since *in vivo* concentrations of patients are currently unknown. We fully acknowledge that this observation is far from firmly revealing the mechanisms explaining the discrepant therapeutic activities of UDCA and NorUDCA in patients^6^, but our findings may at least in part offer interesting mechanistic insights and indicate new pathways by which NorUDCA regulates hepatic inflammation. Therefore future clinical trials for NorUDCA and UDCA should include assessment of *in vivo* concentrations both compounds could achieve and biomarkers or tests evaluating their effects on T cells. Additionally, it raises exciting possibilities that targeted manipulation of bile acid chemical structures (e.g. removal of a single methyl group in NorUDCA^25, 26^) may add on novel signaling and immunomodulatory activities to bile acid and creates additional future therapeutic avenues.

Overall, our findings may have imminent therapeutic implications for treatment of PSC, where NorUDCA has shown promising results ^7^ while UDCA has no established efficacy ^6^. Further studies may be warranted to explore the array of potential therapeutic applications of NorUDCA in immune-mediated liver diseases.

## Materials and Methods

### CD8^+^ T cell Isolation, Activation and Proliferation Assay

CD8^+^ T cells were isolated from peripheral lymph nodes and spleens of C57BL/6J male mice (bred in the mouse facility of Medical University of Vienna) and purified by negative depletion. The purity of the cells was assessed by flow cytometry and was routinely >90%. CD8^+^ T cells were labeled with eBioscience cell proliferation dye eFluor450 as recommended by the manufacturer, stimulated *in vitro* with plate-bound anti-CD3 (1μg/ml) and anti-CD28 (3μg/ml) Ab and expanded in RPMI 1640 medium containing 10% fetal calf serum (FCS), GlutaMAX (2mM, ThermoFisher), β-mercaptoethanol (50μM, ThermoFisher) and Penicillin-Streptomycin (ThermoFisher). To test the impacts of BAs on CD8^+^ T cell activation and proliferation, culture medium was supplemented ± UDCA or NorUDCA for indicated durations and concentrations. Viability of the cells was monitored by a fixable viability dye eFluor 506 (ThermoFisher). Dilution of cell proliferation dye was evaluated by flow cytometry (LSRII Fortessa, BD Biosciences).

### Compound Treatments

Concentrations of NorUDCA (500μM) and UDCA (50μM) for *in vitro* experiments were selected to mimic the *in vivo* serum bile acid level of mice on 0.5% (wt/wt) NorUDCA or UDCA-supplemented diet for 10 days^8^ (supplementary Fig. 3). Concentrations of NorUDCA and UDCA selected for *in vitro* experiments were assessed by viability staining on CD8^+^ T cells. All the other compounds were used at the indicated concentrations: Rapamycin (mTOR inhibitor; 100 nM, Cell signaling), Ly294002 (PI3K inhibitor; 10 μM, Cell signaling), 2-Deoxy-D-glucose (2-DG; 2 mM, Sigma-Aldrich).

### Flow Cytometric Analysis

Murine CD8^+^ T cells cultured for 48h with plate-bound anti-CD3/anti-CD28 Abs were stimulated with 2ng/ml PMA (Sigma-Aldrich) and 1μg/ml ionomycin (Sigma-Aldrich) in the presence of Golgi stop (BD) for cytokine and transcription factor studies, and subjected to intracellular staining using Cytofix/Cytoperm (BD) and permeabilization buffer (BD) for phosphorylation studies. CD8^+^ T cells were fixed 10 min at 37 °C with Cytofix (BD), permeabilized 20 min on ice with methanol and stained with the indicated phospho-Abs for 30 min in the dark at 4°C in PBS/2% FCS. For extracellular staining, CD8^+^ T cells were incubated with Abs for 30 min on ice.

Single cell suspensions of spleen and liver were obtained through mechanical disruption through 70 μm cell strainers. Intrahepatic T cells were enriched following gradient separation^27^. Total cell count was quantified using a Beckman Coulter Counter and normalized to tissue weight. For circulating immune cells assessment, blood was pipetted into Heparin containing tubes and treated with RBC lysis buffer (eBioscience). Samples were then treated with FcR-Block (clone 93, eBioscience), and subsequently stained with the respective antibodies as indicated in each experiment.

Tetramer staining (GP33/GP276/NP396-specific tetramers from NIH Tetramer Core Facility, US) was performed at 37°C for 15min prior to FcR-block treatment.

Murine peripheral blood was treated by Lyse/Fix buffer (BD) for 10min at 37 °C and cells permeabilized with Perm/Wash I buffer (BD) for 30min on ice. Staining with anti-pRPS6 Abs (Cell Signaling) was performed for 30min on ice. Antibodies are listed in Supplementary Table 2.

### Immunoblot Analysis

Immunoblot analysis was performed using standard protocols as previously described using antibodies shown in Supplementary Table 2.

### Phosphatidic Acid (PA) Analysis

Murine CD8^+^ T cells were activated for 24h ± NorUDCA or Rapamycin and harvested for the measurement of intracellular PA using the fluorometric PicoProbe™ PA Assay Kit (Biovision) following manufacturer’s instructions.

### Seahorse Metabolic Assay

Oxygen consumption rates and extracellular acidification rates of activated CD8^+^ T cells ± NorUDCA treatment were measured as previously described^13^.

### Lactate Measurements

Supernatants and cells were collected from 24h murine CD8^+^ T cell culture ± NorUDCA or Rapamycin upon activation and processed following the manufacturer’s instructions (Cayman). Final concentrations of lactate were normalized to cell counts.

### LCMV Clone13 Mouse Model

8 weeks old C57BL/6J male mice (Janvier labs, France) were pre-fed with standard chow or 0.5% (wt/wt) NorUDCA or UDCA-supplemented diet for 10 days or treated with Rapamycin (600 μg/Kg) i.p. 1 day before infection^16, 28^ with 2×10^6^ focus-forming units (FFU) of LCMV strain clone13 intravenously^12^. Mice were continued on NorUDCA or UDCA-containing diet or standard chow or treated with Rapamycin i.p. daily^16, 28^ and sacrificed at day10 following infection, a time point when CD8^+^ T cell inflammation response reaches a peak *in vivo*^10^. Uninfected naive mice on standard chow diet were studied as control. Virus titers were determined by focus-forming assay as previously described Experiments were approved by the Federal Ministry for Science and Research at Medical University of Vienna (BMWFW-66.009/0361-WF/V/3b/2017).

### Ex vivo re-stimulation of splenocytes

Splenocytes from LCMV clone13-infected mice were isolated and incubated with gp33-41 peptide (0.4μg/mL) for 5h at 37 °C in the presence of CD107a/b Ab and stained with the phospho-RPS6 (Cell Signaling) anti-IFN-γ and anti-TNF-α Abs (BD) using Cytofix/Cytoperm (BD) according to manufacturer’s recommendations. Analysis was performed by using FlowJo software (Tree Star). Clones of antibodies used are shown in Supplementary Table 2.

### Histopathology and Multicolor-Immunofluorescence Staining

Fixed liver tissues were embedded in paraffin, sectioned, and stained with immunofluorescent Abs to determine the distribution and frequency of tissue infiltrating CD3^+^CD8^+^ T cells and pRP^S6Ser235/236+^CD3^+^ T cells. 5 representative images (magnification, 400x) were captured from each mouse liver slide for quantitative analysis using ImageJ software V.1.47v (National Institute of Health).

### Serum Analysis

Serum biochemistry was performed as described previously^8^.

### Gene Expression Analysis

RNA isolation from liver, cDNA synthesis and real-time PCR were performed as described previously^8^. Gene expression was calculated relative to *Hprt*. Primer sequences are provided in Supplementary Table 3.

### Human T cell experiments

3 independent experiments with peripheral T cells obtained from 3 PSC patients (also suffering from associated inflammatory bowel disease) and 3 age and gender matched healthy volunteers and were performed following the Declaration of Helsinki and approved by the Ethics Committee of the MUV: 747/2011 and 2001/2018. Bulk T cells from peripheral blood mononuclear cells (PBMCs) were isolated by negative depletion^29^, labeled with 1μM CFSE and rested overnight in RPMI 1640 medium with 5% heat-inactivated FCS, 2mM L-glutamine, 100μg/ml streptomycin and 100U/ml penicillin. T cells were stimulated with plate-bound anti-CD3 (1μg/ml) plus soluble anti-CD28 (0.5μg/ml) Abs ± NorUDCA or UDCA or Rapamycin. Lymphoblastogenesis, proliferation and activity of mTORC1 were assessed on day 3. Effector function of CD8^+^ T cells was assessed on day 6 on a Fortessa flow cytometer (BD Biosciences) and quantified according to the CFSE peaks with FlowJo (Tree Star)^29^. Clones/antibodies are shown in Supplementary Table 2.

### Quantification and statistical analysis

All values are expressed as mean ± standard error of the mean and were statistically analyzed as detailed in the figure legends using GraphPad Prism v.7.0 (La Jolla, CA, USA). A *P* value of less than 0.5 was considered statistically significant and indicated as follows: *= *P*<0.05, **= *P*<0.01, ***= *P*<0.001, ****=*P*<0.0001.

## Acknowledgements

The authors thank Dr. Doreen A. Cantrell and Dr. Jens L Hukelmann from University of Dundee for kindly sharing the detailed protocol of sample processing for proteomics and phosphoproteomics analysis. Our thanks go to Dr. E. John Wherry, Dr. Bertram Bengsch from University of Pennsylvania Perelman School Medicine and Dr. Koichi Araki and Dr. Rafi Ahmed for kindly sharing Rapamycin *in vivo* application protocol.

## Author Contributions

C.Z., N.B., W.E. and M.T. designed the research; C.Z. and N.B. performed most of the experiments and analyzed the data; C.Z., N.B., A.C.M. and P.M. performed proteomic and phosphoproteomic experiments and analyzed the data; C.Z., N.B., H.B., C.V., T.C., C.D.F., A.B., D.H., T.P., L.S., M.A., A.F.G., M. K., P.H., J.R., and S.S. performed *in vivo* experiments; C.Z., N.B., A.O-R., P.S., C.D., E.H. and H.Stockinger. performed *in vitro* human T cell experiments and analyzed the data. C.Z., H.B., H.S. and T.S. measured serum liver biochemistry. C.Z., N.B., A.L., A.G., S.S., L.S., P.H. and T.C. performed metabolic assays. C.Z., N.B., W.E. and M.T. wrote the manuscript. All authors read and corrected the manuscript.

