## Supplementary figures for "24-Nor-Ursodeoxycholic acid reshapes immunometabolism in CD8^+^ T cells and alleviates hepatic inflammation"

### Supplementary Fig.1

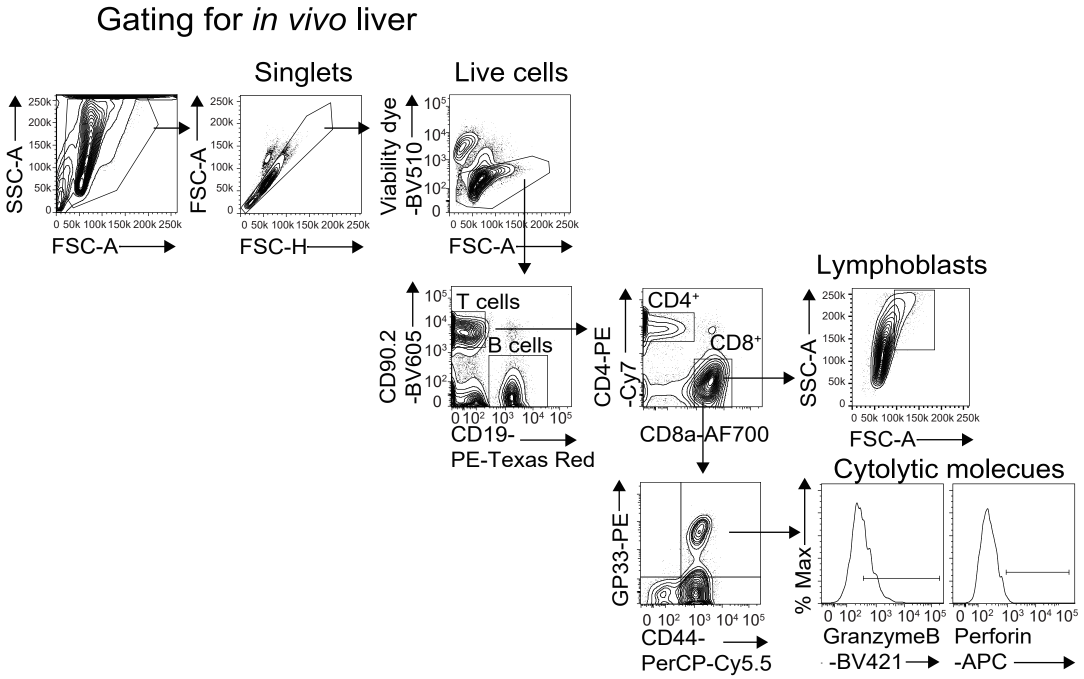

### Supplementary Fig.2

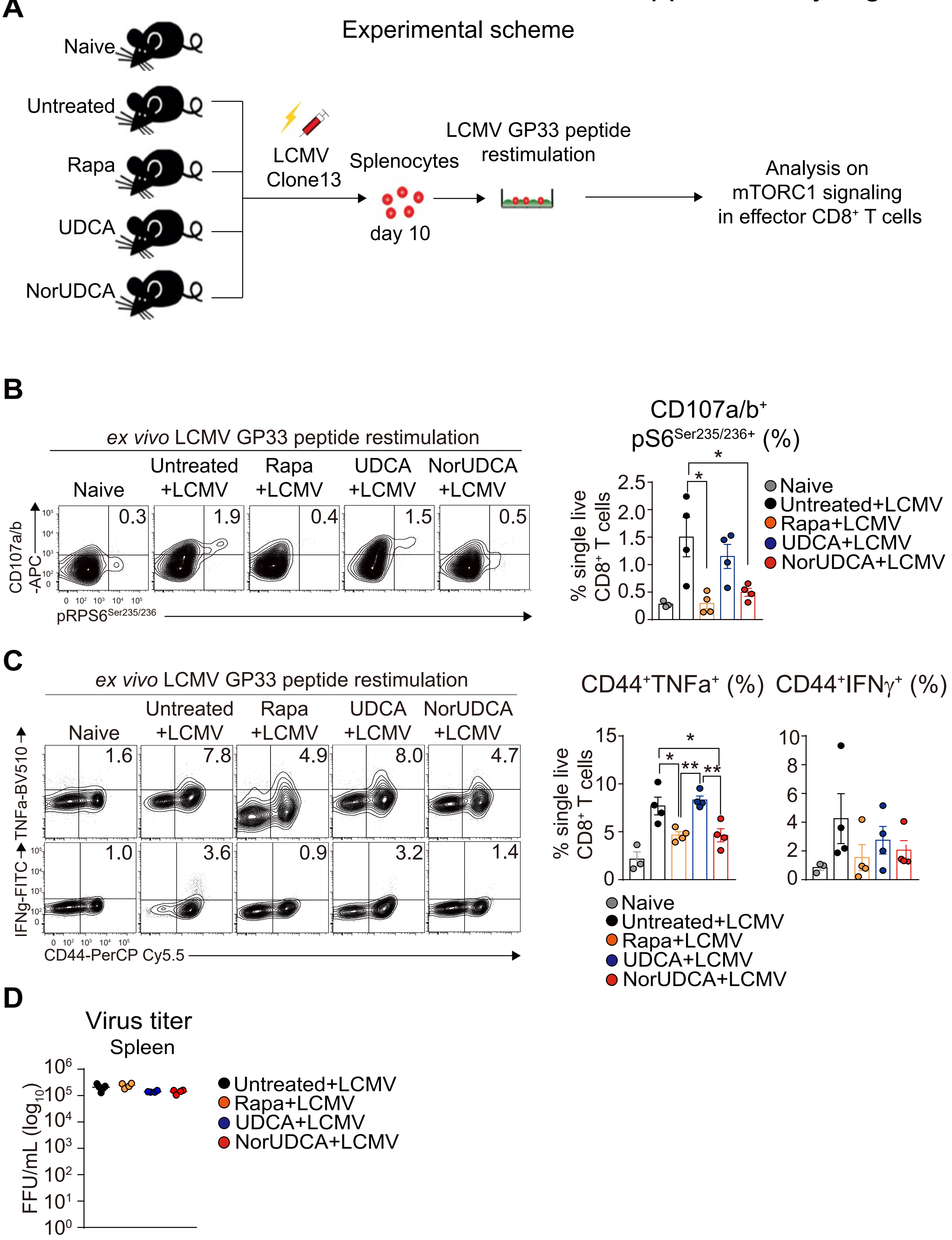

Gating for *in vitro* murine CD8$^+$ T cell culture

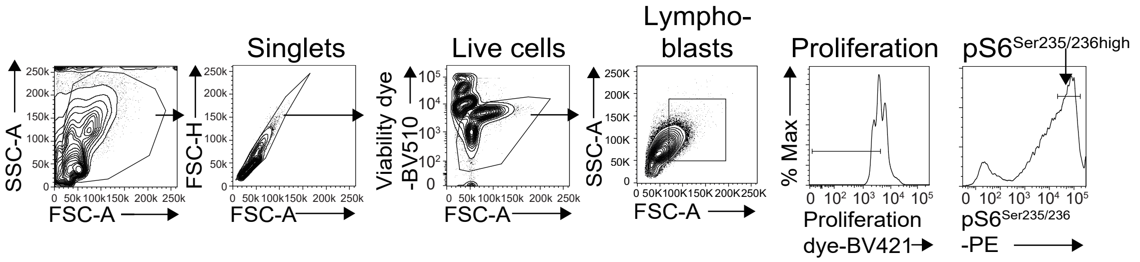

Supplementary Fig.4

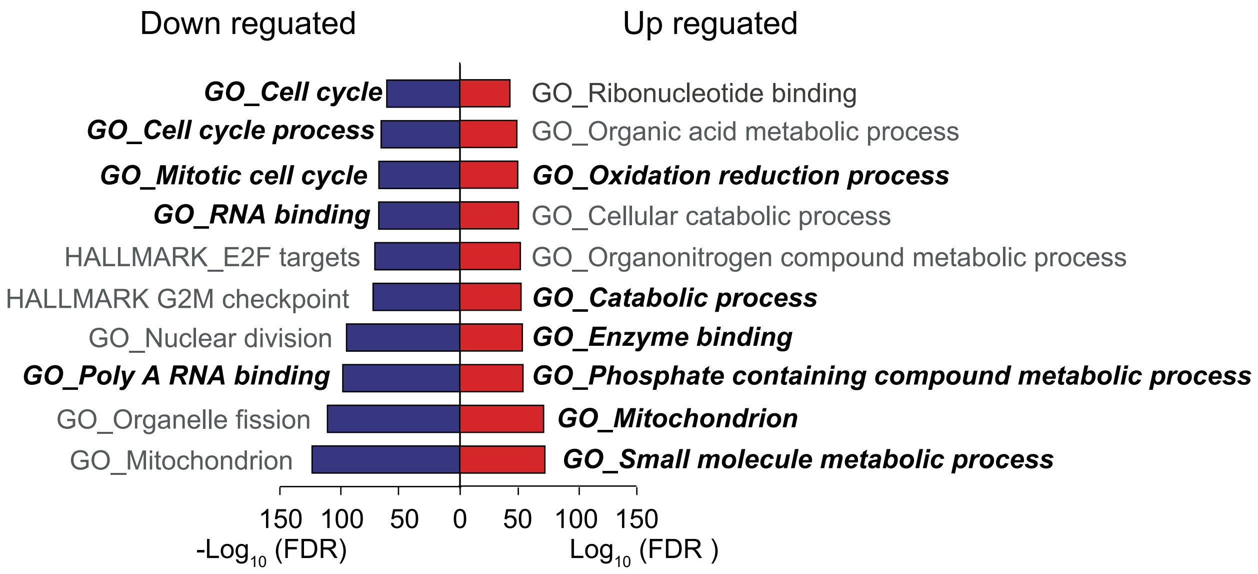

### Supplementary Fig.5

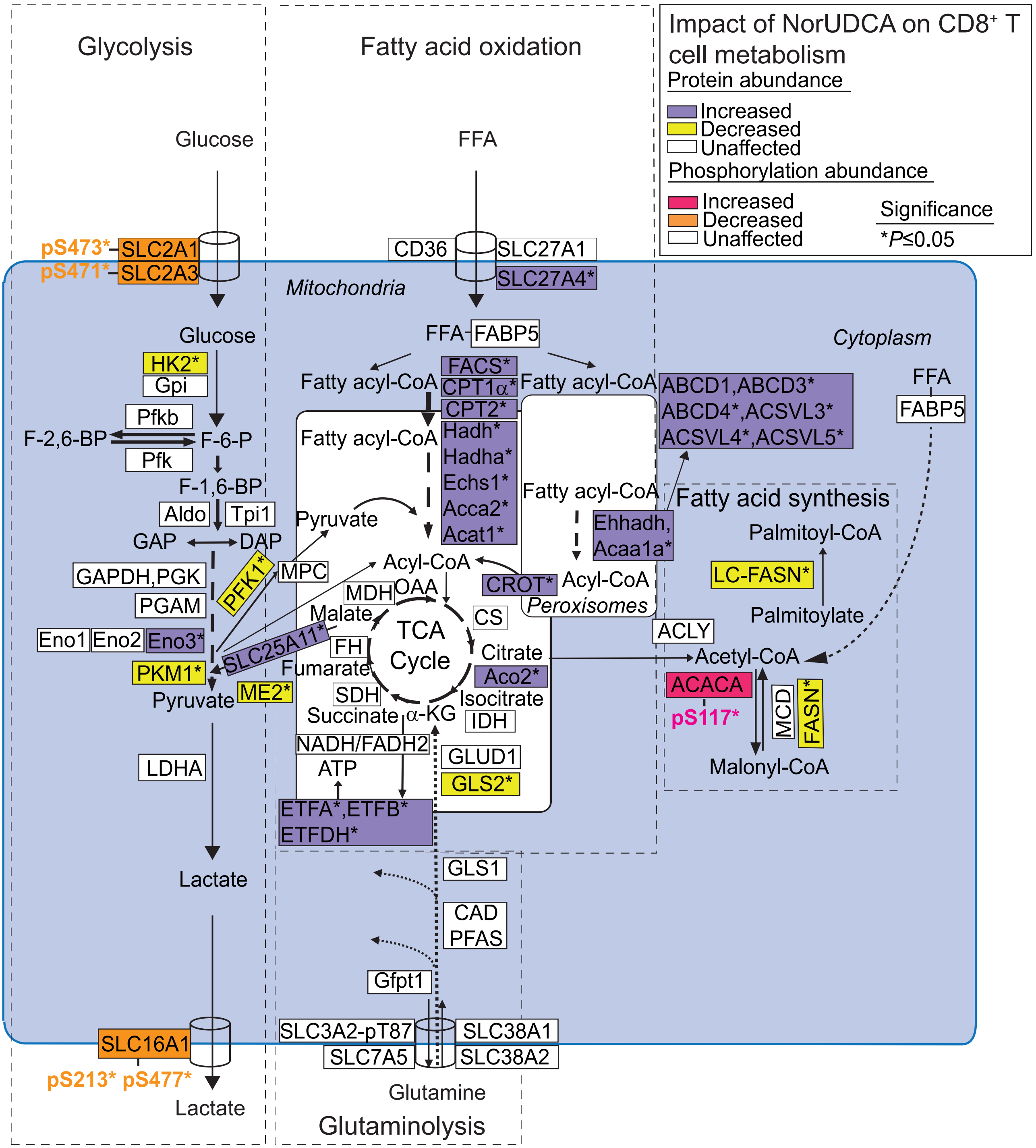

### Supplementary Fig.6

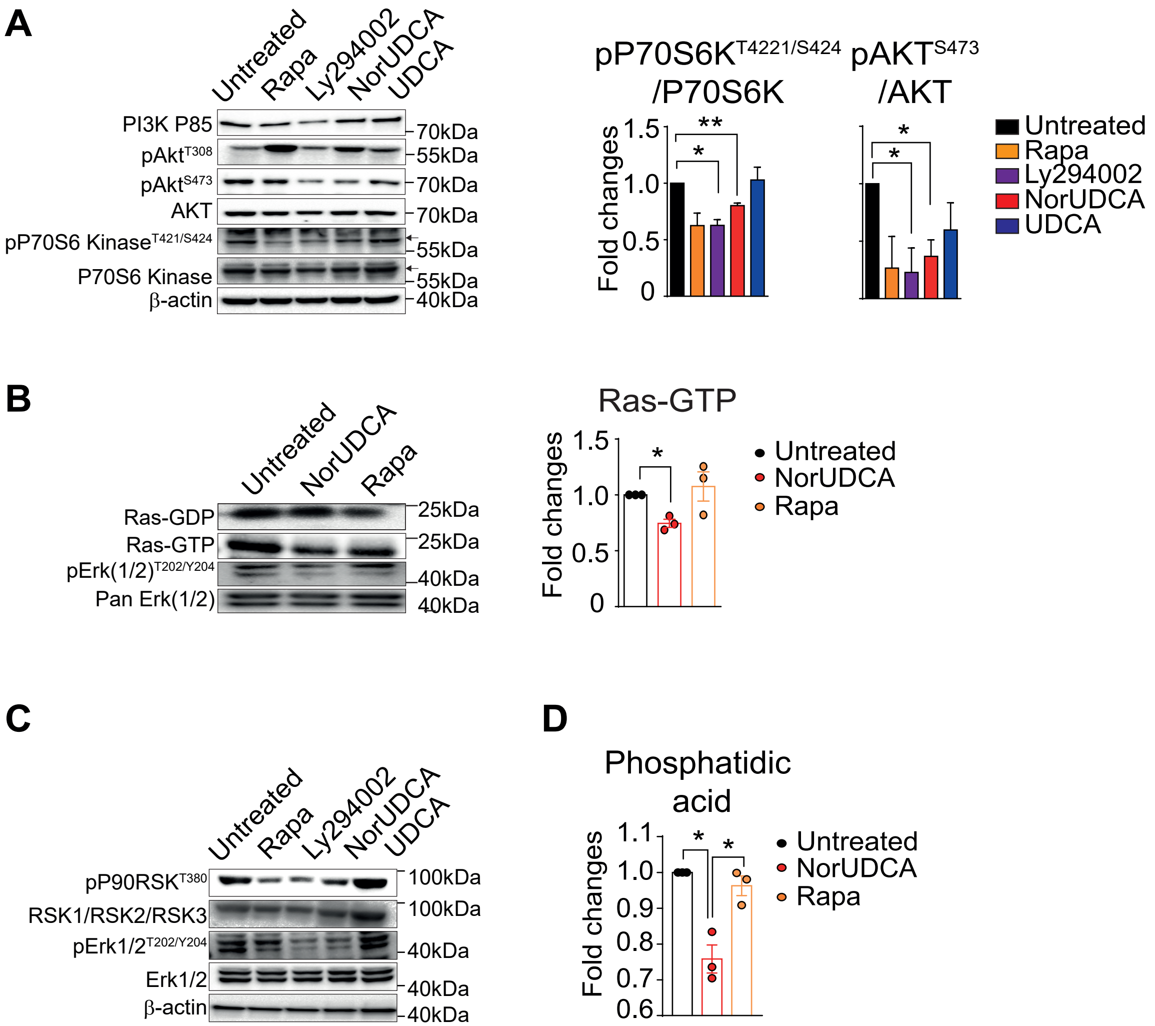

### Supplementary Fig.7

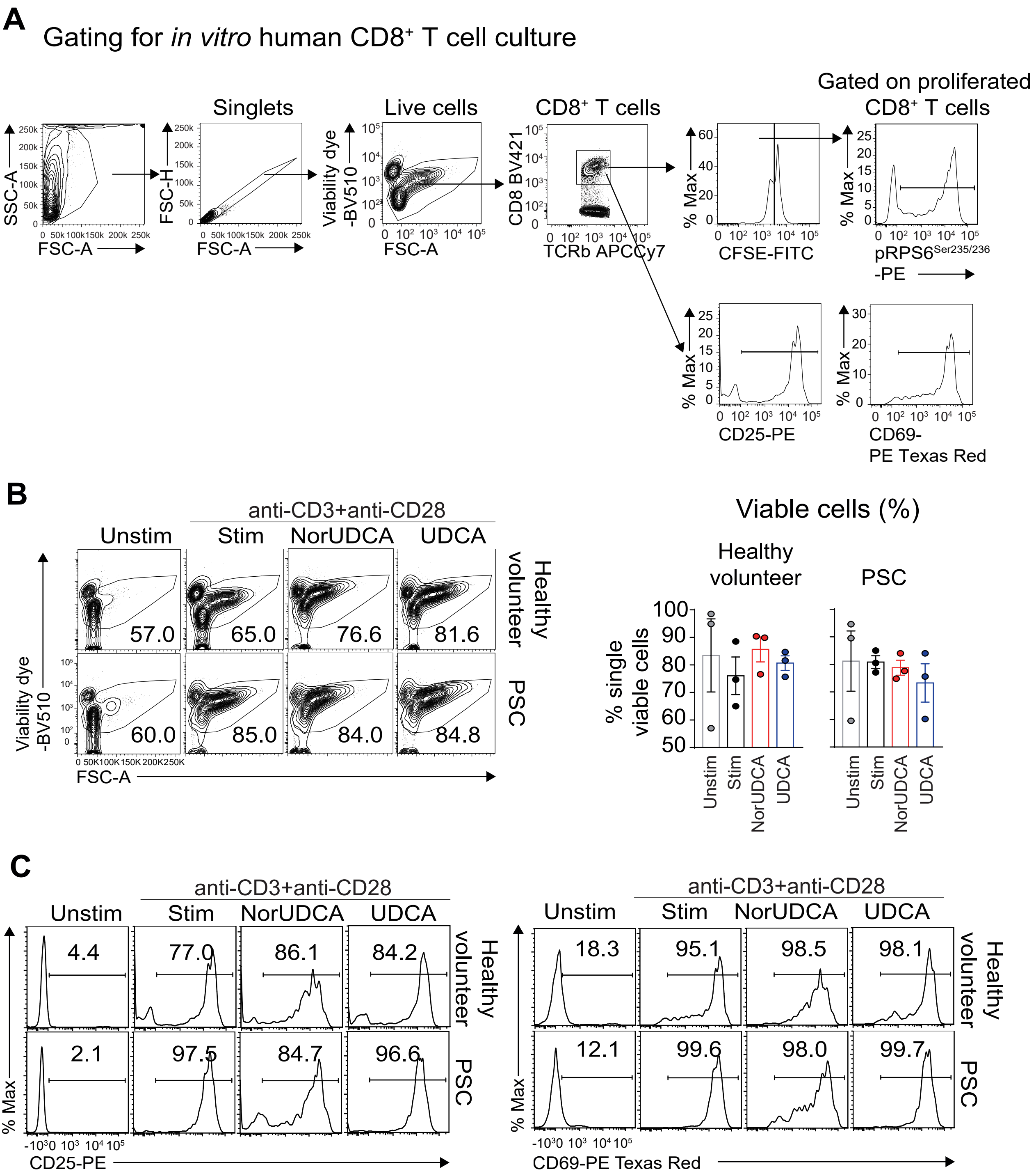
