## Supplementary materials, supplementary figure legends, tables for "24-Nor-Ursodeoxycholic acid reshapes immunometabolism in CD8^+^ T cells and alleviates hepatic inflammation"

**Supplementary Material and Methods**

**Calcium Influx**

Activated CD8^+^ T cells were labeled at 37 °C for 1h with Indo-1 (ThermoFisher), uncoupled Indo-1 was washed out and the cells further incubated ± NorUDCA for 1h at 37 °C. Cells were then washed twice with RPMI medium containing 10% FCS, GlutaMAX, β-mercaptoethanol and antibiotics and resuspended in the same medium ± NorUDCA. Cells were incubated with 2 µg anti-CD3 Ab in 100 µl for 15min at room temperature (RT). Cells were mixed with 400 µl of 37°C pre-warmed RPMI-1640 medium ± NorUDCA, baseline ratio of violet to blue (405nm/510nm) was measured on a BD-LSR II (Becton Dickinson) flow cytometer for 1 min, then 5 µg streptavidin was added to cross-link the TCR. Calcium influx was measured based on the change in the ratio of violet to blue fluorescence for additional 6-7 minutes. Calcium data were analyzed using the FlowJo software.

**Short Term Activation Assay**

As described previously, to induce T cell receptor (TCR) crosslinking, murine splenocytes were freshly isolated, subsequently subjected to 1h pre-incubation with NorUDCA at 37°C and incubated with soluble biotinylated anti-CD3 for 15min at RT. Cells were then brought up to 37°C and linked via addition of streptavidin for indicated duration. In case of human T cells, untouched T cells were pre-incubated with or without NorUDCA at 37°C, loaded with the anti-CD3 and anti-CD28 Ab (5μg/ml each) for 5 min on ice, then brought up to 37°C and cross-linked with sterile goat anti-mouse IgG + IgM Abs (Jackson ImmunoResearch Laboratories, West Grove, PA) for the indicated time points. Reaction was stopped by fixation buffer for 10 minutes. Then cells were washed, stained extracellularly for CD4, CD8, CD19 and CD44 and permeablized for intracellular staining of phospho-antibodies. Antibody clone information is available in Supplementary Table 1.

**Proteomics Sample Preparation and Phosphopeptide Enrichment**

Primary murine CD8^+^ T cells activated with anti-CD3 and anti-CD28 for 24h ± NorUDCA or Rapamycin were harvested in 3 biological replicates for each condition. Each washed cell pellet was lysed separately in 40 μL of freshly prepared lysis buffer containing 50 mM HEPES (pH 8.0), 2% SDS, 0.1 M DTT, 1 mM PMSF, phosSTOP and protease inhibitor cocktail (Sigma-Aldrich). Samples were rested at RT for 20 minutes before heating to 99°C for 5 min. After cooling down to RT, DNA was sheared by sonication using a Covaris S2 high performance ultrasonicator. Cell debris was removed by centrifugation at 20, 000g for 15 min at 20°C. Supernatants were transferred to fresh eppendorf tubes and protein concentration determined using the BCA protein assay kit (Pierce Biotechnology, Rockford, IL). FASP was performed using a 30 kDa molecular weight cutoff filter (VIVACON 500; Sartorius Stedim Biotech GmbH, 37, 070 Goettingen, Germany)^1^. In brief, 43 µg total proteins per sample were reduced by adding DTT at a final concentration of 83.3 mM followed by incubation at 99°C for 5 min. After cooling to room temperature, samples were mixed with 200 μL of freshly prepared 8 M urea in 100 mM Tris-HCl (pH 8.5) (UA-solution) in the filter unit and centrifuged at 14, 000 g for 15 min at 20 °C to remove SDS. Any residual SDS was washed out by a second washing step with 200 μL of UA. The proteins were alkylated with 100 μL of 50 mM iodoacetamide in the dark for 30 min at RT. Afterward, three washing steps with 100 μL of UA solution were performed, followed by three washing steps with 100µL of 50 mM TEAB buffer (Sigma-Aldrich). Proteins were digested with trypsin at a ratio of 1:50 overnight at 37 °C. Peptides were recovered using 40 μL of 50 mM TEAB buffer followed by 50 μL of 0.5 M NaCl (Sigma-Aldrich). Peptides were desalted using C18 solid phase extraction spin columns (The Nest Group, Southborough, MA). After desalting, peptides were labeled with TMT 10plex™ reagents according to the manufacturer (Pierce, Rockford, IL). After quenching of the labeling reaction, labeled peptides were pooled; organic solvent removed in vacuum concentrator and labeled peptides loaded onto a SPE column. Peptides were eluted with 300µL 80% acetonitrile containing 0.1% trifluoroacetic to achieve a final peptide concentration of ~1 µg/µl. Eluate was then used for phosphopeptide enrichment applying a modified method of immobilized metal affinity chromatography (IMAC) ^2^. Briefly, two times 100 µL of Ni-NTA superflow slurry (QIAGEN Inc., Valencia, USA) were washed with LCMS-grade water and Ni2+ stripped off the beads by incubation with 100 mM of EDTA, pH 8 solution for 1h at RT. Stripped NTA resin was recharged with Fe3+ ions by incubation with a fresh solution of Fe(III)Cl3 and 100 µL of charged resin slurry used for the enrichment of a total of ~400 µg TMT-labeled peptides. The unbound fraction was transferred to a fresh glass vial and used for offline fractionation for the analysis of the whole proteome. After washing the slurry with 0.1% TFA, phosphopeptides were eluted with a freshly prepared ammonia solution containing 3mM EDTA, pH 8 and all used for offline fractionation for the analysis of the phophoproteome.

**Offline Fractionation via RP-HPLC at High pH**

Tryptic peptides were re-buffered in 20 mM ammonium formiate buffer shortly before separation by reversed phase liquid chromatography at pH10. The unbound fraction of the phosphopeptide enrichment was separated into 96 time-based fractions on a Phenomenex column (150×2.0 mm Gemini-NX 3µm C18 110Å, Phenomenex, Torrance, CA, USA) using an Agilent 1200 series HPLC system fitted with a binary pump delivering solvent at 100µL/min. Acidified fractions were consolidated into 40 fractions via a concatenated strategy^3^. The bound fraction containing the phosphopeptides was separated into 20 fractions on a Dionex column (500µm × 50mm PepSwift RP, monolithic, Dionex Corporation, Sunnyvale, CA, USA) using an Agilent 1, 200 series nanopump delivering solvent at 4 µL/min. Peptides were separated by applying a gradient of 90% aceonitrile containing 20mM ammonium formiate, pH 10^4^. After solvent removal in a vacuum concentrator, samples were reconstituted in 5% formic acid for LC-MS/MS analysis and kept at -80°C until analysis.

**2D-RP/RP Liquid Chromatography Mass Spectrometry**

Mass spectrometry was performed on an Orbitrap Fusion Lumos mass spectrometer (ThermoFisher Scientific, San Jose, CA) coupled to an Dionex Ultimate 3000RSLC nano system (ThermoFisher Scientific, San Jose, CA) via nanoflex source interface. Tryptic peptides were loaded onto a trap column (Pepmap 100 5μm, 5×0.3 mm, ThermoFisher Scientific, San Jose, CA) at a flow rate of 10μL/min using 2% ACN and 0.1% TFA as loading buffer. After loading, the trap column was switched in-line with a 30 cm, 75 µm inner diameter analytical column (packed in-house with ReproSil-Pur 120 C18-AQ, 3μm, Dr. Maisch, Ammerbuch-Entringen, Germany). Mobile-phase A consisted of 0.4% formic acid in water and mobile-phase B of 0.4% formic acid in a mix of 90% acetonitrile and 10% water. The flow rate was set to 230 nL/min and a 90 min gradient used (6 to 30% solvent B within 81min, 30 to 65% solvent B within 8min and, 65 to 100% solvent B within 1min, 100% solvent B for 6min before equilibrating at 6% solvent B for 18min). Analysis was performed in a data-dependent acquisition mode. Full MS scans were acquired with a scan range of 375-1650 m/z in the orbitrap at a resolution of 120,000 (at 200Th). Automatic gain control (AGC) was set to a target of 2 x 105 and a maximum injection time of 50ms. Precursor ions for MS2 analysis were selected using a TopN dependent scan approach with a max cycle time of 3seconds. MS2 spectra were acquired in the orbitrap (FT) at a resolution of 50,000 (at 200Th). Precursor isolation in the quadrupole was set to 1 Da and higher energy collision induced dissociation (HCD) with normalized collision energy (NCE) of 38%. AGC was set to 5x104 with a maximum injection time of 54ms and 150ms for the proteome and phosphoproteome, respectively. Dynamic exclusion for selected ions was 60s for the proteome and 30s for the phosphoproteome. A single lock mass at m/z 445.120024 for recalibration was employed ^5^. Xcalibur version 4.0.0 and Tune 2.1 were used to operate the instrument. Phosphoproteomics samples were acquired in two technical replicates.

**Proteomics and Phosphoproteomics Data Analysis**

Acquired raw data files were processed using the Proteome Discoverer 2.2.0. platform, utilizing the Sequest HT database search engine and Percolator validation software node (V3.04) to remove false positives with a false discovery rate (FDR) of 1% on peptide and protein level under strict conditions. Searches were performed with full tryptic digestion against the mouse SwissProt database v2017.12 appended with known contaminants (25,293 sequences) with up to two miscleavage sites. Oxidation (+15.9949 Da) of methionine, deamidation (+0.984 Da) of glutamine and asparagine, and protein N-termini acetylation (+42.011 Da) were set as variable modifications, whilst carbamidomethylation (+57.0214 Da) of cysteine residues and TMT 6-plex labeling of peptide N-termini and lysine residues were set as fixed modifications. For phosphopeptides phosphorylation (+79.9663 Da) of serine, threonine and tyrosine was additionally included as a variable modification. Data was searched with mass tolerances of ±10 ppm and 0.025 Da on the precursor and fragment ions (HCD), respectively. Results were filtered to include peptide spectrum matches (PSMs) with Sequest HT cross-correlation factor (Xcorr) scores of ≥1 and 1% FDR peptide confidence. The ptmRS algorithm was additionally used to validate phospopeptides with a set score cutoff of 90. PSMs with precursor isolation interference values of ≥ 50% and average TMT-reporter ion signal-to-noise values (S/N) ≤ 10 were excluded from quantitation. Isotopic impurity correction and TMT channel-normalization based on total peptide amount were applied. Protein abundances were further normalized to equal median in each TMT channel. One-way ANOVA test corrected for false discovery rate by Benjamini–Hochberg procedure followed by Tukey post-hoc test was used to calculate statistical significance of comparisons made among 3 groups of observed changes on protein level. In cases where only the data on NorUDCA and control samples were compared the statistical significance was calculated by paired Student’s t-test. TMT ratios with P-values lower than 0.01 were considered as significant. For the phosphopeptide analysis, peptides that were changing in opposite directions in the two technical replicates were discarded, moreover Student’s t-test was performed on each technical replicate separately and P-value was required to be lower than 0.01 in both replicates in order to consider phosphopeptide abundance change as significant. The mean abundance of each biological replicate from the two technical replicates was calculated and further normalized to their corresponding proteome abundance.

**Supplementary Figure legends**

**Supplementary Fig. 1. Gating strategy for *in vivo* LCMV experiment.** Gating strategy for flow cytometry analysis of *in vivo* LCMV experiment.

**Supplementary Fig. 2. Splenic CD8^+^ T cells from NorUDCA treated LCMV mice show reduced mTORC1 and ameliorated TNFα expression but unaltered IFNγ expression upon *ex vivo* re-stimulation with GP33-LCMV peptide.** (A) Experimental scheme. (B) CD44 or pRPS6^Ser235/236^ expression in *ex vivo* isolated splenic CD107a/b^+^CD8^+^ T cells upon GP33 LCMV peptide re-stimulation. Quantitative analysis is shown alongside. (C) TNFα or IFNγ expression on gated live singlet splenic CD107a/b^+^CD8^+^ T cells isolated from indicated groups upon *ex vivo* LCMV GP33 peptides re-stimulation. Quantitative analysis is shown alongside. Data is representative of 2 independent experiments. At least 3 biological independent animals were used per group during experiment. Summary data are presented as mean±SE. *P* values were calculated by one-way ANOVA corrected with Tukey post-hoc test. *=*P*<0.05, **=*P*<0.01, ***=*P*<0.001, ****=*P*<0.0001. Rapa, Rapamycin.

**Supplementary Fig. 3. Gating strategy for *in vitro* murine CD8^+^ T cell culture.** Gating strategy for flow cytometry analysis of *in vitro* murine CD8^+^ T cell culture.

**Supplementary Fig. 4. The top 10 enriched pathways for significantly up- and downregulated proteins by NorUDCA in CD8^+^ T cells.** The top 10 enriched Hallmark, Gene Ontology (GO), KEGG pathways for significantly up- and downregulated proteins by NorUDCA. Overlap of top 10 enriched pathways modulated by NorUDCA and Rapamycin are shown in bold, black and italic. Pathways uniquely modulated by NorUDCA are shown in grey, regular.

**Supplementary Fig. 5. NorUDCA reduces glycolysis while enhancing fatty acid oxidation machinery in activated CD8^+^ T cells.** Schematic summary of key phosphoproteomics data showing the impact of NorUDCA on glycolysis and fatty acid oxidation pathways. Figure shows protein, protein phosphorylation and genes affected by NorUDCA treatment. The color code was inserted for direction of changes. Data were obtained from 3 independent experiments. *P* values were calculated with two tailed, paired student’s t test between control and NorUDCA groups. *=*P*<0.05.

**Supplementary Fig. 6. NorUDCA attenuates murine Ras-Erk-P90RSK-mTORC1 signaling and reduces intracellular PA level in proliferating CD8^+^ T cells.**

(A,B) Representative immunoblots of Ras-Erk-P90RSK-mTORC1 pathway in activated CD8^+^ T cells. Quantitative analysis is shown alongside. (B) Intracellular PA of CD8^+^ T cells stimulated under indicated conditions. Data are representative of 3 independent experiments. Quantitative data are presented as mean±SE. *P* values were calculated by one-way ANOVA corrected with Tukey post-hoc test. Data are normalized to protein abundance of untreated group. *=*P*<0.05, **=*P*<0.01. Unstim, unstimulated, Stim,stimulated, Rapa, Rapamycin.

**Supplementary Fig. 7. NorUDCA does not affect human CD8^+^ T cell viability and activation.** (A) Gating strategy for flow cytometry analysis of in vitro human CD8^+^ T cell culture. (B) Viability profile of circulating CD8^+^ T cells from peripheral blood of healthy volunteer and PSC patients treated under indicated for 3 days. Plots show one representative experiments. Quantitative analysis of 3 independent experiments is shown alongside. (C) Expression of CD25 and CD69 on live singlet CD8^+^ T cells of healthy volunteer and PSC patient are shown. Data in (B,C) are representative 3 independent experiments. *P* values were calculated by one-way ANOVA corrected with Tukey post-hoc test. PSC, primary sclerosing cholangitis; Rapa, Rapamycin.

**Supplementary Table 1. Proteins involved in the GO_Mitochondrion pathways upregulated and downregulated by NorUDCA.**

| Proteins modulated by NorUDCA in downregulation of Mitochorion pathways | |
| --- | --- |
| Protein name |  |
| TP53 | tumor protein p53 |
| DNA2 | DNA replication helicase/nuclease 2 |
| AKAP8 | A-kinase anchoring protein 8 |
| PNPT1 | polyribonucleotide nucleotidyltransferase 1 |
| TYMS | thymidylate synthetase |
| DHFR | dihydrofolate reductase |
| CDK1 | cyclin dependent kinase 1 |
| DLGAP5 | DLG associated protein 5 |
| RAD51 | RAD51 recombinase |
| MSTO1 | misato mitochondrial distribution and morphology regulator 1 |
| TFDP1 | transcription factor Dp-1 |
| GADD45GIP1 | GADD45G interacting protein 1 |
| PIN1 | "peptidylprolyl cis/trans isomerase, NIMA-interacting 1 |
| HAUS3 | HAUS augmin like complex subunit 3 |
| NME6 | NME/NM23 nucleoside diphosphate kinase 6 |
| AKT1 | AKT serine/threonine kinase 1 |
| GADD45GIP1 | GADD45G interacting protein 1 |
| PIN1 | "peptidylprolyl cis/trans isomerase, NIMA-interacting 1 |
| HAUS3 | HAUS augmin like complex subunit 3 |
| NME6 | NME/NM23 nucleoside diphosphate kinase 6 |
| AKT1 | AKT serine/threonine kinase 1 |
| RPS3 | ribosomal protein S3 |
| TOP3A | DNA topoisomerase III alpha |
| SLC25A33 | solute carrier family 25 member 33 |
| MYO19 | myosin XIX |
| MRPL41 | mitochondrial ribosomal protein L41 |
| HJURP | Holliday junction recognition protein |
| CASP8AP2 | Caspase 8 assciate protein 2 |
| CDK5RAP1 | CDK5 regulatory subunit associated protein 1 |
| CRY1 | cryptochrome circadian regulator 1 |
| APEX2 | apurinic/apyrimidinic endodeoxyribonuclease 2 |
| DAP3 | death associated protein 3 |
| NSUN4 | NOP2/Sun RNA methyltransferase 4 |
| MTERF4 | mitochondrial transcription termination factor 4 |
| TRMT10C | tRNA methyltransferase 10C, mitochondrial RNase P subunit |
| MRPS9 | mitochondrial ribosomal protein S9 |
| MRPL40 | mitochondrial ribosomal protein L40 |
| MRPL11 | mitochondrial ribosomal protein L11 |
| MRPL22 | mitochondrial ribosomal protein L22 |
| MRPS7 | mitochondrial ribosomal protein S7 |
| MRPL58 | mitochondrial ribosomal protein L58 |
| MRPL9 | mitochondrial ribosomal protein L9 |
| MRPL13 | mitochondrial ribosomal protein L13 |
| MRPS18A | mitochondrial ribosomal protein S18A |
| MRPL15 | mitochondrial ribosomal protein L15 |
| MRPL3 | mitochondrial ribosomal protein L3 |
| MRPL28 | mitochondrial ribosomal protein L28 |
| MRPL54 | mitochondrial ribosomal protein L54 |
| MRPL2 | mitochondrial ribosomal protein L2 |
| MRPL4 | mitochondrial ribosomal protein L4 |
| MRPL37 | mitochondrial ribosomal protein L37 |
| MRPL27 | mitochondrial ribosomal protein L27 |
| MRPL32 | mitochondrial ribosomal protein L32 |
| MRPL45 | mitochondrial ribosomal protein L45 |
| MRPS21 | mitochondrial ribosomal protein S21 |
| MRPS14 | mitochondrial ribosomal protein S14 |
| MRPS5 | mitochondrial ribosomal protein S5 |
| MRPL1 | mitochondrial ribosomal protein L1 |
| YARS2 | tyrosyl-tRNA synthetase 2 |
| AARS2 | alanyl-tRNA synthetase 2, mitochondrial |
| ERAL1 | Era like 12S mitochondrial rRNA chaperone 1 |
| GFM2 | G elongation factor mitochondrial 2 |
| TUFM | Tu translation elongation factor, mitochondrial |
| GFM1 | G elongation factor mitochondrial 1 |
| RCC1L | RCC1 like |
| NOA1 | nitric oxide associated 1 |
| MTIF2 | mitochondrial translational initiation factor 2 |
| TRUB2 | TruB pseudouridine synthase family member 2 |
| FASTKD2 | FAST kinase domains 2 |
| NGRN | neugrin, neurite outgrowth associated |
| PTCD3 | pentatricopeptide repeat domain 3 |
| MTIF3 | mitochondrial translational initiation factor 3 |
| TEFM | transcription elongation factor, mitochondrial |
| FASTKD5 | FAST kinase domains 5 |
| POLRMT | RNA polymerase mitochondrial |
| CD3EAP | CD3e molecule associated protein |
| HSPD1 | heat shock protein family D (Hsp60) member 1 |
| LGALS3 | galectin 3 |
| SLIRP | SRA stem-loop interacting RNA binding protein |
| KARS1 | lysyl-tRNA synthetase 1 |
| FARS2 | "phenylalanyl-tRNA synthetase 2, mitochondrial |
| IREB2 | iron responsive element binding protein 2 |
| ASS1 | argininosuccinate synthase 1 |
| DDX28 | DEAD-box helicase 28 |
| TRMU | tRNA 5-methylaminomethyl-2-thiouridylate methyltransferase |
| MTPAP | mitochondrial poly(A) polymerase\ |
| MRM3 | mitochondrial rRNA methyltransferase 3 |
| GRSF1 | G-rich RNA sequence binding factor 1 |
| PTCD1 | pentatricopeptide repeat domain 1 |
| KYAT3 | kynurenine aminotransferase 3 |
| PTCD2 | pentatricopeptide repeat domain 2 |
| AKAP1 | A-kinase anchoring protein 1 |
| MAPK8 | mitogen-activated protein kinase 8 |
| DCTPP1 | dCTP pyrophosphatase 1 |
| MRPL19 | mitochondrial ribosomal protein L19 |
| MRPL17 | mitochondrial ribosomal protein L17 |
| MRPL47 | mitochondrial ribosomal protein L47 |
| MRPL57 | mitochondrial ribosomal protein L57 |
| MRPL24 | mitochondrial ribosomal protein L24 |
| MRPL33 | mitochondrial ribosomal protein L33 |
| MRPL53 | mitochondrial ribosomal protein L53 |
| MRPL48 | mitochondrial ribosomal protein L48 |
| MRPL38 | mitochondrial ribosomal protein L38 |
| MRPS22 | mitochondrial ribosomal protein S22 |
| MRPS33 | mitochondrial ribosomal protein S33 |
| TARS2 | "threonyl-tRNA synthetase 2, mitochondrial |
| HARS1 | histidyl-tRNA synthetase 1 |
| QRSL1 | glutaminyl-tRNA amidotransferase subunit QRSL1 |
| MTG1 | mitochondrial ribosome associated GTPase 1 |
| MTERF3 | mitochondrial transcription termination factor 3 |
| DHODH | dihydroorotate dehydrogenase (quinone) |
| BNIP3 | BCL2 interacting protein 3 |
| MTFR2 | mitochondrial fission regulator 2 |
| GSK3A | glycogen synthase kinase 3 alpha |
| GLRX2 | glutaredoxin 2 |
| TIGAR | TP53 induced glycolysis regulatory phosphatase |
| MARS2 | methionyl-tRNA synthetase 2, mitochondrial |
| METTL17 | methyltransferase like 17 |
| THOP1 | thimet oligopeptidase 1 |
| NLN | neurolysin |
| FPGS | folylpolyglutamate synthase |
| PDHB | pyruvate dehydrogenase E1 beta subunit |
| DEGS1 | delta 4-desaturase, sphingolipid 1 |
| NMNAT3 | nicotinamide nucleotide adenylyltransferase 3 |
| KIF1B | kinesin family member 1B |
| RAF1 | Raf-1 proto-oncogene, serine/threonine kinase |
| STING1 | stimulator of interferon response cGAMP interactor 1 |
| PI4KB | phosphatidylinositol 4-kinase beta |
| RIPK3 | receptor interacting serine/threonine kinase 3 |
| SPATA5 | spermatogenesis associated 5 |
| RIPK2 | receptor interacting serine/threonine kinase 2 |
| MMAA | metabolism of cobalamin associated A |
| GTPBP3 | GTP binding protein 3, mitochondrial |
| PPIF | peptidylprolyl isomerase F |
| SDHAF2 | succinate dehydrogenase complex assembly factor 2 |
| GLS | glutaminase |
| TMLHE | trimethyllysine hydroxylase, epsilon |
| NDUFB8 | NADH:ubiquinone oxidoreductase subunit B8 |
| NUDT9 | nudix hydrolase 9 |
| COQ5 | coenzyme Q5, methyltransferase |
| NDUFAF7 | NADH:ubiquinone oxidoreductase complex assembly factor 7 |
| PDSS2 | decaprenyl diphosphate synthase subunit 2 |
| MPST | mercaptopyruvate sulfurtransferase |
| PITRM1 | pitrilysin metallopeptidase 1 |
| ME2 | malic enzyme 2 |
| ALDH1B1 | aldehyde dehydrogenase 1 family member B1 |
| GCAT | glycine C-acetyltransferase |
| QTRT1 | queuine tRNA-ribosyltransferase catalytic subunit 1 |
| RBFA | ribosome binding factor A |
| COX7A2L | cytochrome c oxidase subunit 7A2 like |
| CIAPIN1 | cytokine induced apoptosis inhibitor 1 |
| ACAT2 | acetyl-CoA acetyltransferase 2 |
| TAMM41 | TAM41 mitochondrial translocator assembly and maintenance homolog |
| MT-CO2 | mitochondrially encoded cytochrome c oxidase II |
| SLC25A13 | solute carrier family 25 member 13 |
| BLOC1S2 | biogenesis of lysosomal organelles complex 1 subunit 2 |
| FOXRED1 | FAD dependent oxidoreductase domain containing 1 |
| NDUFAF2 | NADH:ubiquinone oxidoreductase complex assembly factor 2 |
| PISD | phosphatidylserine decarboxylase |
| L2HGDH | L-2-hydroxyglutarate dehydrogenase |
| COQ4 | coenzyme Q4 |
| GHITM | growth hormone inducible transmembrane protein |
| CHCHD2 | coiled-coil-helix-coiled-coil-helix domain containing 2 |
| PGAM5 | "PGAM family member 5, mitochondrial serine/threonine protein phosphatas |
| FMC1 | formation of mitochondrial complex V assembly factor 1 homolog |
| PTRH2 | peptidyl-tRNA hydrolase 2 |
| GZMB | granzyme B |
| BSG | basigin (Ok blood group) |
| SACS | sacsin molecular chaperone |
| KIFBP | kinesin family binding protein |

| Proteins modulated by NorUDCA in upregulation of Mitochorion pathways | |
| --- | --- |
| Protein name | Discription |
| PKM | pyruvate kinase M1/2 |
| IDH1 | isocitrate dehydrogenase (NADP(+)) 1 |
| CYB5R3 | cytochrome b5 reductase 3 |
| CAT | catalase |
| ARSB | arylsulfatase B |
| GSTP1 | glutathione S-transferase pi 1 |
| PARK7 | Parkinsonism associated deglycase |
| CPT1A | carnitine palmitoyltransferase 1A |
| GLUD1 | glutamate dehydrogenase 1 |
| MECP2 | methyl-CpG binding protein 2 |
| HADH | hydroxyacyl-CoA dehydrogenase |
| IDH2 | isocitrate dehydrogenase (NADP(+)) 2 |
| GLUL | glutamate-ammonia ligase |
| ACSL4 | acyl-CoA synthetase long chain family member 4 |
| OXCT1 | 3-oxoacid CoA-transferase 1 |
| ABCD3 | ATP binding cassette subfamily D member 3 |
| ALDH18A1 | aldehyde dehydrogenase 18 family member A1 |
| ADH5 | "alcohol dehydrogenase 5 (class III), chi polypeptide |
| ACAT1 | acetyl-CoA acetyltransferase 1 |
| PYCR1 | pyrroline-5-carboxylate reductase 1 |
| SLC25A12 | solute carrier family 25 member 12 |
| HADHB | hydroxyacyl-CoA dehydrogenase trifunctional multienzyme complex subunit beta |
| SCP2 | sterol carrier protein 2 |
| ACADL | acyl-CoA dehydrogenase long chain |
| HADHA | hydroxyacyl-CoA dehydrogenase trifunctional multienzyme complex subunit alpha |
| ACSS1 | acyl-CoA synthetase short chain family member 1 |
| SUCLG2 | succinate-CoA ligase GDP-forming beta subunit |
| SUCLA2 | succinate-CoA ligase ADP-forming beta subunit |
| ECH1 | enoyl-CoA hydratase 1 |
| ACAA2 | acetyl-CoA acyltransferase 2 |
| HSD17B10 | hydroxysteroid 17-beta dehydrogenase 10 |
| IVD | isovaleryl-CoA dehydrogenase |
| ETFA | electron transfer flavoprotein subunit alpha |
| ACADS | acyl-CoA dehydrogenase short chain |
| ECHS1 | enoyl-CoA hydratase, short chain 1 |
| ECI1 | enoyl-CoA delta isomerase 1 |
| ALDH7A1 | aldehyde dehydrogenase 7 family member A1 |
| MTHFD2 | methylenetetrahydrofolate dehydrogenase (NADP+ dependent) 2, methenyltetrahydrofolate cyclohydrolase |
| SUCLG1 | succinate-CoA ligase alpha subunit |
| COQ9 | coenzyme Q9 |
| FXN | frataxin |
| GARS1 | glycyl-tRNA synthetase 1 |
| PON2 | paraoxonase 2 |
| GSTZ1 | glutathione S-transferase zeta 1 |
| MMUT | methylmalonyl-CoA mutase |
| KYAT1 | kynurenine aminotransferase 1 |
| RIDA | reactive intermediate imine deaminase A homolog |
| ACSL5 | acyl-CoA synthetase long chain family member 5 |
| GNPAT | glyceronephosphate O-acyltransferase |
| PCCA | propionyl-CoA carboxylase subunit alpha |
| PCCB | propionyl-CoA carboxylase subunit beta |
| PCK2 | phosphoenolpyruvate carboxykinase 2, mitochondrial |
| AARS1 | alanyl-tRNA synthetase 1 |
| MCCC2 | methylcrotonoyl-CoA carboxylase 2 |
| MTHFS | methenyltetrahydrofolate synthetase |
| EARS2 | "glutamyl-tRNA synthetase 2, mitochondrial |
| HIBCH | 3-hydroxyisobutyryl-CoA hydrolase |
| PPA2 | inorganic pyrophosphatase 2 |
| PDE2A | phosphodiesterase 2A |
| PARP1 | poly(ADP-ribose) polymerase 1 |
| NAXE | NAD(P)HX epimerase |
| AGPAT4 | 1-acylglycerol-3-phosphate O-acyltransferase 4 |
| CNP | "2',3'-cyclic nucleotide 3' phosphodiesterase |
| PNKP | polynucleotide kinase 3'-phosphatase |
| DTYMK | deoxythymidylate kinase |
| DGUOK | deoxyguanosine kinase |
| AK3 | adenylate kinase 3 |
| MMAB | metabolism of cobalamin associated B |
| FOXO1 | forkhead box O1 |
| ATP5PO | ATP synthase peripheral stalk subunit OSCP |
| ATP5PD | ATP synthase peripheral stalk subunit d |
| CLYBL | citrate lyase beta like |
| ISCA2 | iron-sulfur cluster assembly 2 |
| SLC25A1 | solute carrier family 25 member 1 |
| SFXN3 | sideroflexin 3 |
| PYCARD | PYD and CARD domain containing |
| NCSTN | nicastrin |
| CAPN1 | calpain 1 |
| PPP3CA | protein phosphatase 3 catalytic subunit alpha |
| CCR7 | C-C motif chemokine receptor 7 |
| FYN | "FYN proto-oncogene, Src family tyrosine kinase |
| BCL2 | BCL2 apoptosis regulator |
| CASP8 | caspase 8 |
| STAP1 | signal transducing adaptor family member 1 |
| DNAJA3 | DnaJ heat shock protein family (Hsp40) member A3 |
| SRI | sorcin |
| ARL2 | ADP ribosylation factor like GTPase 2 |
| SLC25A4 | solute carrier family 25 member 4 |
| IDE | insulin degrading enzyme |
| SOD2 | superoxide dismutase 2 |
| RTN4IP1 | reticulon 4 interacting protein 1 |
| NDUFA13 | NADH:ubiquinone oxidoreductase subunit A13 |
| TRAP1 | TNF receptor associated protein 1 |
| NDUFV1 | NADH:ubiquinone oxidoreductase core subunit V1 |
| GSTK1 | glutathione S-transferase kappa 1 |
| WWOX | WW domain containing oxidoreductase |
| PRDX5 | peroxiredoxin 5 |
| CYCS | cytochrome c, somatic |
| NDUFA8 | NADH:ubiquinone oxidoreductase subunit A8 |
| NDUFS8 | NADH:ubiquinone oxidoreductase core subunit S8 |
| NDUFV3 | NADH:ubiquinone oxidoreductase subunit V3 |
| NDUFA6 | NADH:ubiquinone oxidoreductase subunit A6 |
| NDUFB5 | NADH:ubiquinone oxidoreductase subunit B5 |
| NDUFB10 | NADH:ubiquinone oxidoreductase subunit B10 |
| COX15 | cytochrome c oxidase assembly homolog COX15 |
| UQCRC1 | ubiquinol-cytochrome c reductase core protein 1 |
| HSDL2 | hydroxysteroid dehydrogenase like 2 |
| ALKBH3 | alkB homolog 3, alpha-ketoglutaratedependent dioxygenase |
| FAHD1 | fumarylacetoacetate hydrolase domain containing 1 |
| ANXA6 | annexin A6 |
| SQSTM1 | sequestosome 1 |
| ATP5IF1 | ATP synthase inhibitory factor subunit 1 |
| ALAS1 | 5'-aminolevulinate synthase 1 |
| LACTB | lactamase beta |
| HSCB | HscB mitochondrial iron-sulfur cluster cochaperone |
| PRKACA | protein kinase cAMP-activated catalytic subunit alpha |
| UBA1 | ubiquitin like modifier activating enzyme 1 |
| ABCB6 | ATP binding cassette subfamily B member 6 (Langereis blood group) |
| MAP2K2 | mitogen-activated protein kinase kinase 2 |
| NAPG | NSF attachment protein gamma |
| RPS27A | ribosomal protein S27a |
| BCL2L11 | BCL2 like 11 |
| CAPN2 | calpain 2 |
| PNPLA7 | patatin like phospholipase domain containing 7 |
| FOXO3 | forkhead box O3 |
| NIPSNAP2 | nipsnap homolog 2 |
| VDAC3 | voltage dependent anion channel 3 |
| DNAJC11 | DnaJ heat shock protein family (Hsp40) member C11 |
| ACBD3 | acyl-CoA binding domain containing 3 |
| ABCB8 | ATP binding cassette subfamily B member 8 |
| GIMAP8 | GTPase, IMAP family member 8 |
| HCLS1 | hematopoietic cell-specific Lyn substrate 1 |
| PAM16 | presequence translocase associated motor 16 |
| PPP3CC | protein phosphatase 3 catalytic subunit gamma |
| MTCH2 | mitochondrial carrier 2 |
| DNAJC19 | DnaJ heat shock protein family (Hsp40) member C19 |
| PMPCA | peptidase, mitochondrial processing alpha subunit |
| PARL | presenilin associated rhomboid like |
| TIMM8B | translocase of inner mitochondrial membrane 8 homolog B |
| ACP6 | acid phosphatase 6, lysophosphatidic |
| HDHD5 | haloacid dehalogenase like hydrolase domain containing 5 |
| GRAMD4 | GRAM domain containing 4 |
| FAM162A | family with sequence similarity 162 member A |
| URI1 | URI1 prefoldin like chaperone |
| SLC25A24 | solute carrier family 25 member 24 |
| SLC25A20 | solute carrier family 25 member 20 |
| TFAM | "transcription factor A, mitochondrial |
| COA3 | cytochrome c oxidase assembly factor 3 |
| COX16 | cytochrome c oxidase assembly factor COX16 |
| FECH | ferrochelatase |
| TBRG4 | transforming growth factor beta regulator 4 |
| LACTB2 | lactamase beta 2 |
| POLDIP2 | DNA polymerase delta interacting protein 2 |
| DCAF8 | DDB1 and CUL4 associated factor 8 |
| SLC25A39 | solute carrier family 25 member 39 |
| CLIC4 | chloride intracellular channel 4 |
| SLC25A45 | solute carrier family 25 member 45 |
| SFXN2 | sideroflexin 2 |
| CHCHD5 | coiled-coil-helix-coiled-coil-helix domain containing 5 |
| KLC2 | kinesin light chain 2 |
| ARMC10 | armadillo repeat containing 10 |
| GLOD4 | glyoxalase domain containing 4 |
| CISD3 | CDGSH iron sulfur domain 3 |
| AKAP10 | A-kinase anchoring protein 10 |
| BOLA3 | bolA family member 3 |
| NIPSNAP3B | nipsnap homolog 3B |

**Supplementary Table 2. Antibodies used in the study.**

| **Antigen** | **Clone** | **Company** | **Application** |
| --- | --- | --- | --- |
| CD4 | RM4-5 | Biolegend | F.C. |
| CD8a | 53-6.7 | Biolegend | F.C. |
| CD11b | M1/70 | Biolegend | F.C. |
| CD11c | N418 | Biolegend | F.C. |
| CD45R | B220; RA3-6B2 | Biolegend | F.C. |
| Gr1 | RB6-8C5 | Biolegend | F.C. |
| Ter-119 | TER-119 | Biolegend | F.C. |
| NK1.1 | PK136 | Biolegend | F.C. |
| CD25 | PC61 | Biolegend | F.C. |
| CD69 | H1.2F3 | Biolegend | F.C. |
| CD62L | MEL-14 | Biolegend | F.C. |
| CD8a | SK1 | Biolegend | F.C. |
| CD4 | SK3 | Biolegend | F.C. |
| CD69 | FN50 | Biolegend | F.C. |
| CD25 | BC96 | Biolegend | F.C. |
| Perforin | eBioOMAK-D | Thermofisher | F.C. |
| TCRa/b | IP26 | Biolegend | F.C. |
| IL-2 | JES6-5H4 | BD Biosciences | F.C. |
| Granzyme B | GB11 | BD Biosciences | F.C. |
| INFγ | XMG1.2 | BD Biosciences | F.C. |
| TNFα | MP6-XT22 | BD Biosciences | F.C. |
| T-bet | 4B10 | BD Biosciences | F.C. |
| CD3 | 145-2C11 | BD Biosciences | F.C. |
| CD28 | 37.51 | BD Biosciences | F.C. |
| CD3 | OKT3 | BD Biosciences | F.C. |
| CD28 | CD28.2 | BD Biosciences | F.C. |
| GLUT1 | EPR3915 | Abcam | F.C. |
| CD3 | SP7 | Novus | I.F. |
| CD8 | 53-6.7 | Novus | I.F. |
| Rabbit IgG | Polyclonal | Invitrogen | I.F. |
| pmTOR(Ser2448) | D9C2 | Cell Signalling Technology | W.B. |
| mTOR | Polyclonal | Cell Signalling Technology | W.B. |
| pP70S6 Kinase (T421/S424) | Polyclonal | Cell Signalling Technology | W.B. |
| P70S6 Kinase | Polyclonal | Cell Signalling Technology | W.B. |
| pAKT(Ser473) | Polyclonal | Cell Signalling Technology | W.B. |
| AKT | Polyclonal | Cell Signalling Technology | W.B. |
| Beta-actin | 8H10D10 | Cell Signalling Technology | W.B. |
| pPI3 Kinase p85 (Tyr458)/p55 (Tyr199) | Polyclonal | Cell Signalling Technology | W.B. |
| PI3 Kinase p85 | Polyclonal | Cell Signalling Technology | W.B. |
| PI3 Kinase p55 | D2B3 | Cell Signalling Technology | W.B. |
| PI3K class III PI3 Kinase Class III | D4E2 | Cell Signalling Technology | W.B. |
| PI3 Kinase p110γ | D55D5 | Cell Signalling Technology | W.B. |
| PTEN | D4.3 | Cell Signalling Technology | W.B. |
| pAkt (Thr308) | D25E6 | Cell Signalling Technology | W.B. |
| pP90RSK(T380) | D3H11 | Cell Signalling Technology | W.B. |
| RSK1/RSK2/RSK3 | 32D7 | Cell Signalling Technology | W.B. |
| pP44/42 MAPK(Erk1/2)(T202/Y204) | E10 | Cell Signalling Technology | W.B. |
| P44/42 MAPK(Erk1/2) | Polyclonal | Cell Signalling Technology | W.B. |
| pAMPKα(T172) | D4D6D | Cell Signalling Technology | W.B. |
| AMPKα | D5A2 | Cell Signalling Technology | W.B. |
| CCR7 | 4B12 | Biolegend | F.C. |
| Perforin | S16009A | Biolegend | F.C. |
| CD107a | 1D4B | Biolegend | F.C. |
| CD107b | M3184 | Biolegend |  |
| CD44 | IM7 | Biolegend | F.C. |
| CD19 | 6D5 | Biolegend | F.C. |
| CD90.2 | 30-H12 | Biolegend | F.C. |
| FOXP3 | FJK-16s | Thermofisher | F.C. |
| PD-1 | 29F.1A12 | Biolegend | F.C. |
| CxCR5 | L138D7 | Biolegend | F.C. |
| pRPS6^S235/236^ | D57.2.2E | Cell Signalling Technology | F.C. |
| pRPS6^S240/244^ | D68F8 | Cell Signalling Technology | F.C. |
| pP44/42 MAPK(Erk1/2)(T202/Y204) | E10 | Cell Signalling Technology | F.C. |

Table showing the antibodies used in this study. F.C.: Flow Cytometry W.B.: Western Blot I.F. Immunofluorescence

**Supplementary Table 3. Primer sequences used in the study**

| **Gene** | **Forward** (5’ - 3’) | **Reverse** (5’ - 3’) |
| --- | --- | --- |
| qRTPCR | | |
| *Cd8a* | AGCCAGTGCTGCGAACTCCCT | TCGGCTCCTGTGGTAGCAGATGA |
| *Hprt* | GATACAGGCCAGACTTTGTTG | GGTAGGCTGGCCTATAGGCT |
| *Granzyme b* | ACTCTTGACGCTGGGACCTA | AGTGGGGCTTGACTTCATGT |
| *Tnfα* | GTAGCCCACGTCGTAGCAAAC | AGTTGGTTGTCTTTGAGATCCATG |
| *Cxcl10* | GAATCCGGAATCTAAGACCATCAA | GTGCGTGGCTTCACTCCAGT |
| *Slc2a1 (Glut1)* | AAGTCCTTTGAGATGCTGATCC | CCCACATACATGGGCAC |
| *Hk2* | GGCTAGGAGCTACCACACAC | AACTCGCCATGTTCTGTCCC |
| *Slc16a3* | GACGAGTAGGCTGGACTGAA | GCTGCTTTCACCAAGAACTGA |
| *Ldha* | ATGAGTAAGTCCTCAGGCGG | GGACTTTGAATCTTTTGAGACCTTG |
| *Cpt1a* | TGAGTGGCGTCCTCTTTGG | CAGCGAGTAGCGCATAGTCA |
| *Ppara* | TATTCGGCTGAAGCTGGTGTAC | CTGGCATTTGTTCCGGTTCT |
| *Pparb/d* | AGATGGTGGCAGAGCTATGACC | TCTCCTCCTGTGGCTGTTCC |
| *Pparg1* | TGTGAGACCAACAGCCTGAC | CCGCTTCTTTCAAATCTTGTCTGT |
| *Cd36* | GATCGGAACTGTGGGCTCAT | GGTTCCTTCTTCAAGGACAACTTC |
| *Fabp5* | ACCGAGAGCACAGTGAAGAC | GCCATCAGCTGTCGTTTCATC |
| *Klf2* | ACTTGCAGCTACACCAACTG | GGCTTCTCACCTGTGTGTG |

Table showing the primers used in this study.
